# A thermodynamic framework for mapping elastic recoil mechanism across the human proteome

**DOI:** 10.64898/2026.08.28.747957

**Authors:** Rhea Desai, David Pople, Shreya Jain, Atharwa Musale, Ifrah Sajjad, Richard Wittebort, Ronald L. Koder, Vikas Nanda

**Affiliations:** Center for Advanced Biotechnology and Medicine and the Department of Biochemistry and Molecular Biology, Robert Wood Johnson Medical School, Rutgers, The State University of New Jersey, Piscataway, NJ 08854; Department of Chemistry, University of Louisville, Louisville, KY 40292; Department of Physics, The City College of New York, New York, NY 10031, Graduate Programs of Physics, Chemistry, Biology and Biochemistry, The Graduate Center City University of New York (CUNY), New York, NY 10016

## Abstract

The folding thermodynamics of proteins are dominated by two opposing forces, the loss in backbone entropy and the packing of hydrophobic groups. The same forces are major contributors to the extension thermodynamics of elastic proteins with the distinction that both processes act in concert, favoring the higher chain and solvent entropy of a relaxed conformation. The relative entropic contributions specify the recoil mechanism; human elastin recoil is primarily driven by hydrophobic forces, whereas fly resilin has a rubber-like mechanism driven by backbone entropy. Despite the importance of elastic proteins to tissue biomechanics, few have been identified, let alone characterized to the same extent as elastin and resilin. We develop a thermodynamic framework that maps proteins by sequence-derived estimates of extension-induced backbone and solvent entropy changes. Putative elastic proteins are proposed and classified by recoil mechanism based on estimated thermodynamic features. Proteins that map to elastic regions are overrepresented by the skin proteome. The set of predicted elastic domains is further extended by incorporating sequence context embedded in protein language models. Protein domains with distinct thermodynamic recoil mechanisms cluster on the latent space manifold. Some of these domains are anticipated to have roles within molecular machines, expanding the scope of elastic protein function beyond mechanical materials like elastin and resilin.

**SIGNIFICANCE STATEMENT:** Elastic proteins enable tissues and molecular assemblies to store and recover mechanical energy, yet only a handful, such as elastin and resilin, have been characterized in detail. We introduce a sequence-derived thermodynamic framework that maps proteins according to the relative contributions of backbone conformational entropy and solvent entropy to elastic recoil. Applied to the human proteome, this approach identifies numerous candidate elastic proteins and domains enriched in skin, extracellular matrix, cytoskeletal, and macromolecular assembly functions. Integration with protein language models further reveals that proteins sharing similar recoil mechanisms form distinct neighborhoods in latent space despite limited sequence homology. These findings suggest that elastic function is far more widespread than currently recognized and provide a general strategy for discovering and mechanistically classifying elastic proteins across biological systems.

## INTRODUCTION

Protein folding into a compact, native state is largely driven by hydrophobic forces and opposed by loss of chain degrees of freedom, requiring protein sequences where roughly half of the amino acids are non-polar [1]. The sequence-structure-function paradigm governs about half of the human proteome [2, 3]. The remaining proteins function in a regime of intrinsic disorder. The sequence of an intrinsically disordered protein (IDP) is not coupled to a target native state, but rather to its behavior in a dynamic conformational ensemble. The entropy of an IDP polypeptide chain is a thermodynamic feature determined by sequence and modulated by post translational modifications or conformational selection upon binding specific partners. IDPs play unique biological roles as signaling hubs or interaction scaffolds. The amino acid compositions of IDPs are distinct from folded proteins, with an abundance of polar and charged amino acids, proline and glycine, and a depletion of non-polar amino acids. These lead to unique sequence grammars where composition and non-random sequence patterns can be used to identify general IDP functional classes [4].

An important functional class of IDPs operates not through signaling nor molecular recognition, but through biomechanical elasticity. Elastic proteins are required throughout biology to store mechanical energy, buffer transient forces, and enable reversible deformation in systems ranging from extracellular matrices and connective tissues to molecular machines and cytoskeletal assemblies. In some IDPs, extension reduces the entropy of highly flexible polypeptide chain and surrounding water, generating a restoring force upon release, as is the case with proteins such as elastin and resilin. Elasticity can also arise from the reversible deformation or unfolding of folded domains, as exemplified by proteins such as titin, where mechanical work is stored in structured elements and recovered during refolding [5]. Despite the pervasive need for elastic function in biological systems, remarkably few elastic proteins have been identified and characterized in molecular detail. This may reflect challenges associated with studying elastic proteins, which are often repetitive, intrinsically disordered, crosslinked into insoluble materials or within large supramolecular assemblies. These features complicate structural, biochemical, and genetic analyses, suggesting that elastic proteins may be substantially underrepresented in current functional annotations.

Elastin is the principal elastic protein of vertebrate connective tissues, providing resilience to arteries, lung, skin, and other organs that undergo repeated cycles of deformation [6]. Unlike many mechanical proteins that rely on the unfolding and refolding of structured domains, elastin functions largely as a crosslinked network of intrinsically disordered chains. Its sequence is organized into alternating hydrophobic and lysine-rich domains, where the latter form covalent crosslinks that stabilize the elastic fiber while the former undergo reversible extension and recoil. While classical models proposed that elastin elasticity arose from the loss of configurational entropy of the protein chain upon extension, our current understanding is that recoil is dominated by the hydrophobic effect: extension exposes hydrophobic side chains to solvent, increasing the ordering of surrounding water molecules, while relaxation restores a more compact disordered ensemble and releases these constraints on solvent motion [7]. In this view, elastin behaves as an unusual biological spring in which the principal restoring force is generated by solvent entropy rather than by refolding into a defined native structure.

Another well studied elastic protein is resilin, characterized most extensively in the wing hinges of *Drosophila*, but also where it functions in jump mechanisms, leg joints, feeding pumps, stingers, and acoustic organs across Arthropoda [8]. Unlike elastin, resilin is thought to derive its elasticity primarily from the conformational entropy of the polypeptide chain itself, making its behavior closely analogous to that of natural rubber. Resilin is composed of highly disordered, glycine-rich repetitive sequences that remain mobile even when incorporated into a covalently crosslinked network through di-and tri-tyrosine linkages [9, 10]. Mechanical extension reduces the number of accessible backbone conformations available to these flexible chains, creating an entropic restoring force that drives recoil when the load is removed. Resilin exhibits exceptionally high resilience and can withstand hundreds of millions of loading cycles over an organism’s lifetime. Despite decades of biomechanical study, relatively few resilin-family proteins have been characterized at the molecular level to the same extent as that of *Drosophila* resilin, leaving open the possibility that many proteins with analogous entropic recoil mechanisms remain unidentified.

We explore whether elastic proteins can be identified and classified using a sequence-derived thermodynamic framework that estimates the relative contributions of backbone and solvent entropy to elastic recoil. Applying this framework to the human proteome reveals that proteins with known elastic function, including elastin and resilin, map to discrete regions enriched in intrinsically disordered and mechanically-associated sequences. This approach identifies candidate elastic domains in structural, extracellular matrix, epidermal, and molecular machine proteins that likely would not have been detected through sequence homology alone. While amino acid composition captures major thermodynamic features associated with elasticity, incorporation of sequence grammar through protein language models extends the predictive power of the framework, enabling discrimination between proteins with similar compositions but potentially distinct recoil mechanisms. Together, these results support the view that elastic proteins represent an evolutionarily-diverse and under-annotated functional class within the human proteome, and that thermodynamic features embedded in primary sequence can be used to uncover their molecular recoil mechanisms.

## RESULTS AND DISCUSSION

Sequence homology is insufficient for identifying other elastin-or resilin-like proteins in the human proteome. Human elastin has no clear paralog; when assessed by a PSI-BLAST homology search [11] against the human reference proteome, elastin returns only itself and one other protein, periaxin (with a modest E-value of 0.001). Periaxin is hypothesized to play a mechanical role in myelin sheath integrity and spacing [12] suggesting that it might share broadly-defined elastic function with elastin, but its sequence consists of pentapeptide repeats (VPEMK) that differ physiochemically from elastin pentapeptide repeats (VPGVG), suggesting disparate recoil mechanisms. Fly resilin has no predicted human orthologs as detected using PSI-BLAST. Identifying other putative elastic proteins in the human proteome requires a different approach.

### Thermodynamic Framework

The elastic recoil of resilin is driven by a similar mechanism as natural rubber [13–15], where extension reduces conformational degrees of freedom of a polymer chain (**Fig. 1A**). This decrease in backbone entropy opposes extension, just as it would the folding of a protein into a native state. The higher the sequence content of flexible amino acids such as glycine and alanine, the greater the anticipated loss in backbone entropy upon extension and greater entropic restoring force. An approach is needed to compute the change in backbone entropy between a disordered, relaxed ensemble and a constrained, extended state.

**Figure 1.**
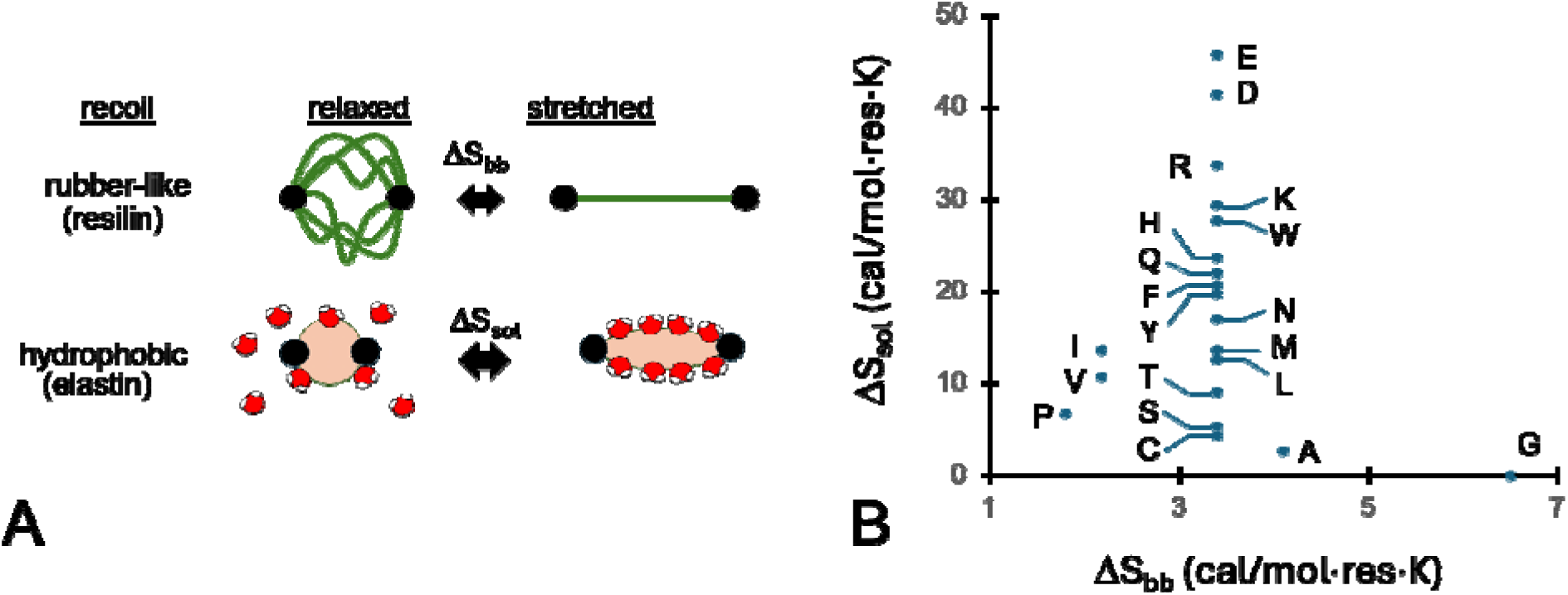
A sequence-based thermodynamic framework for elastic recoil mechanisms. **(A)** Elastic recoil of elastin and resilin are driven by different entropic mechanisms. Loss of chain degrees of freedom upon extension dominates resilin recoil, similar to the elasticity of rubber. Extending elastin increases exposed hydrophobic surfaces, driving recoil by solvent entropy. **(B)** Sequence-based entropy framework where amino acids are mapped based on change in backbone flexibility ΔS_bb_ and solvation entropy ΔS_sol_ upon extension. Parameters listed in **Table S1**.

To estimate the loss in backbone entropy upon extension, we adapt a scale developed by D’Aquino and colleagues [16], where the backbone conformational entropies of amino acids are calculated from the distribution of accessible conformations in /ψ torsional (Ramachandran) space, using N-acetyl-(X-aminoacyl)-N′-methylamide as a model. For the most flexible amino acid, glycine, the calculated ΔS_bb_ is large, 6.5 cal/mol·K, as compared to β-branched amino acids valine and isoleucine (ΔS_bb_ = 2.18 cal/mol·K). Given the model chemical system used, D’Aquino et. al. did not include ΔS_bb_ for proline in their published parameter set. If we assume the fraction of Ramachandran space proline can occupy (F_Pro_) is one-tenth that of glycine [17], we can estimate ΔS_bb_(Pro-Gly) = R·log(F_Pro_/F_Gly_) = -4.7 cal/mol·K. Therefore, ΔS_bb_(Pro) would be 6.5 – 4.7 = 1.8 cal/mol·K. Backbone entropy parameters estimated from NMR order parameters and atomistic molecular simulations [18] are on par with the values used here. These parameters represent an upper-limit on the loss of entropy, assuming maximal extension such that all degrees of freedom are frozen [19].

Elastin differs from resilin in that recoil is driven largely by entropic contributions from water, rather than the protein chain (**Fig. 1A**) [7, 20]. In the relaxed state, hydrophobic sidechains form compact, but disordered ensembles that sequester them from bulk water [21]. Extension exposes these hydrophobic groups, restricting the motion of surface waters and the release of these waters provides a restorative elastic force. The greater the hydrophobicity of the sequence, the larger this restorative force is expected to be. To estimate the magnitude of this effect, we need to compute the change in water entropy when amino acids are exposed upon extension.

The two mechanisms are not mutually exclusive, and all intrinsically disordered or unfolded proteins exhibit some degree of entropic elasticity. The dominant recoil mechanism strongly influences mechanical behavior: solvent-driven recoil can generate restoring forces even at relatively low extensions, whereas entropic recoil is weak at modest extensions and increases sharply as the chain approaches its contour length. The predominance of a rubber-like recoil mechanism in resilin is associated with substantially greater extensibility than is observed for elastin, with resilin capable of strains approaching 300% compared with roughly 50-70% for elastin [22].

To estimate the entropic component of solvent interactions with amino acids, we adopted the scale of Schauperl and colleagues [23]. The entropy of solvation was estimated directly from molecular dynamics trajectories of N-acetyl-(X-aminoacyl)-N′-methylamide, the same chemical model used for calculating backbone entropy. Schauperl analyzed water density and orientation distributions using Grid Inhomogeneous Solvation Theory [24]. Hydration entropies from the TIP3P simulations reported by Schauperl et al. were selected because this water model showed good agreement with experimental hydrophobicity data. In their work, the choice of water model primarily affects the absolute magnitude of the hydration entropy while preserving the relative ordering among amino acids. Notably, while ΔS_bb_ parameters from D’Aquino vary by ∼3 cal/mol·K, the range of ΔS_sol_ is ten-fold larger. The residue-dependent contribution of solvent ordering can be comparable to or larger than the residue-dependent backbone conformational entropy because a single amino acid influences the translational and orientational freedom of many surrounding water molecules, whereas backbone entropy reflects only the conformational freedom of the peptide chain itself. In folding, total conformational entropy change of an amino acid more closely matches changes in solvation entropy [25, 26], but in the case of extension, we expect the entropy associated with backbone degrees of freedom to dominate for proteins with rubber-like recoil mechanisms.

Combining the ΔS_bb_ and ΔS_sol_ scales creates a framework for comparing putative recoil mechanisms across sequences of different proteins (**Fig. 1B**). The backbone component is largely specified by the fractions of proline and glycine residues, which has previously been found to map mechanical features of elastic proteins [27]. The magnitude of solvent entropy depends on a number of factors, including sidechain size, charge and hydrophobicity. We hypothesize proteins with rubber-like elastic recoil mechanisms will be found in the lower right of this map, where ΔS_bb_ is high and ΔS_sol_ is low. Hydrophobicity-driven elastic proteins are expected to map closer to the center of this plot. Given that only a handful of elastic proteins have been characterized, a sequence-based approach such as this could be a powerful framework for identifying novel elastic proteins and anticipating their recoil mechanism.

### Entropy Map of the Human Proteome

In order to place canonical elastic proteins (human elastin and fly resilin) in a broader sequence context, we analyzed the human reference proteome [28], selected because humans are organisms with diverse, well-characterized mechanical tissues, and we hypothesize there are novel elastic proteins to be found in the background distribution of human sequences. Models of the human proteome in AlphaFold DB [29] provide near-complete structural coverage, including confidence estimates that predict distinction between ordered and intrinsically disordered regions at proteome scale [30]. Proteins longer than 1400 residues are split into multiple entries in the AlphaFold DB, resulting in a total of 23,587 entries.

To construct an entropy map, sequence amino acid composition of each entry was converted into average backbone and solvation entropy terms using the D’Aquino and Schauperl-derived scales respectively (**Fig. 2A**). At this stage, calculations were performed over entire protein sequences and therefore average over possible compositional heterogeneity between domains. ΔS_bb_ exhibits a normal distribution, centered at 3.5 cal/mol·K, which is unsurprising given that most amino acids branch at the γ-carbon and share this parameter value. This is also within the uncertainty of thermal energy (k_b_T) of experimental measurements of maximal per-residue backbone entropy loss upon extension [19]. ΔS_sol_ exhibits a normal distribution, centered at 18 cal/mol·K. When combined, the multivariate distribution is normal, but also shows peripheral features trending toward glycine, proline or charge-rich sequences.

**Figure 2.**
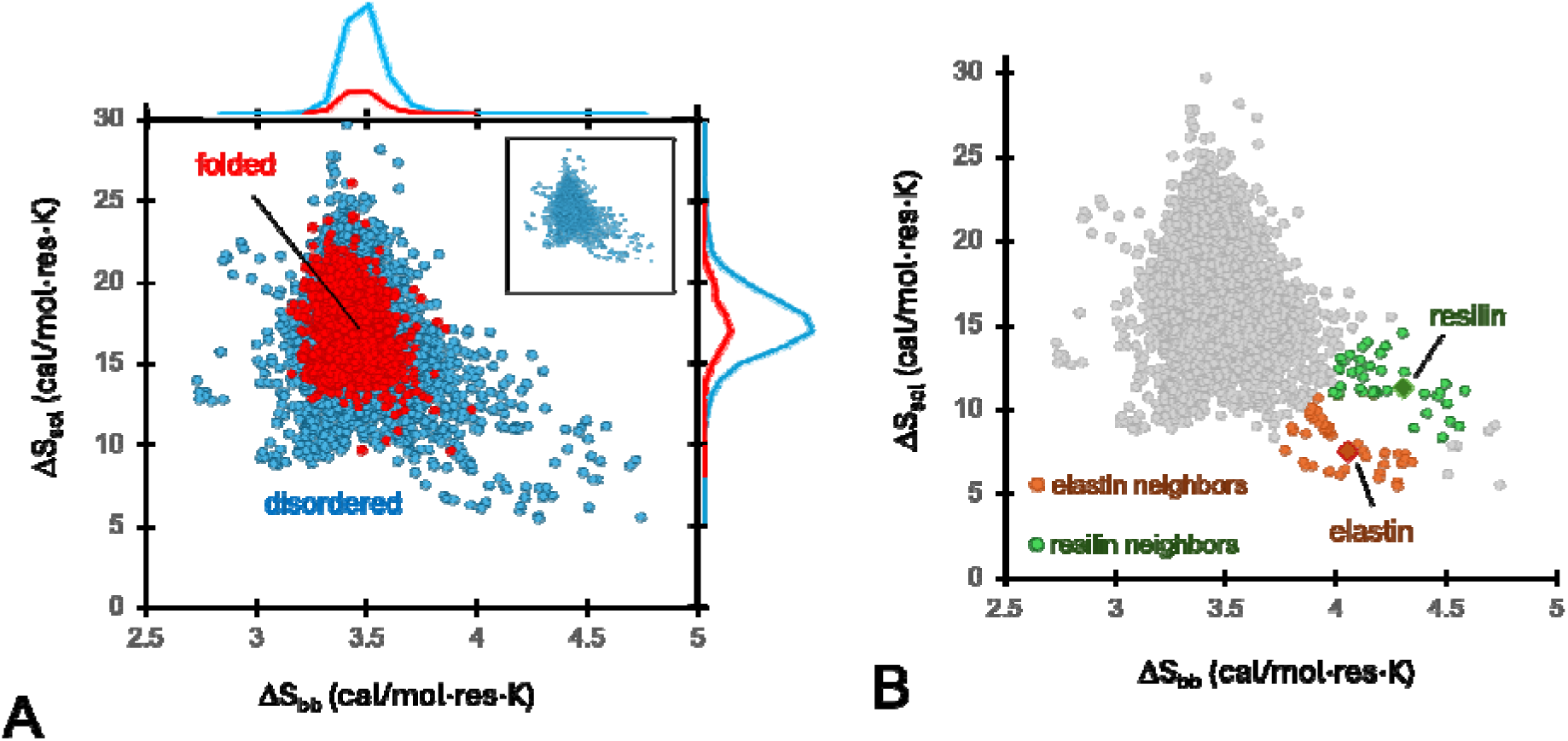
Entropy map of the human proteome. **(A)** Per-residue backbone and solvent entropies for the AlphaFold DB human reference proteome. Folded proteins are defined as entries where >90% of residues have both a pLDDT score > 0.7 and DSSP-specified secondary structural state other than coil. With this strict definition, there are 4048 ordered and 19539 disordered entries. **(inset)** entropy map of disordered entries only. Top and right margins display distributions of solvation and backbone entropies for both folded and disordered entries. **(B)** Euclidean nearest neighbors of human elastin and fly resilin on the entropy map.

Folding into a compact native state requires favorable hydrophobic interactions to balance chain entropy. Entries were classified as folded or disordered based on confidence score cutoffs and DSSP annotations of the associated AlphaFold DB structure models. We used a very stringent definition for ordered proteins, as much of the human proteome contains proteins with both folded domains and intrinsically disordered regions. Ordered proteins had narrower distributions of backbone and solvent entropy and clustered at the center of the plot. Disordered proteins had broader ΔS_bb_ and ΔS_sol_ distributions, consistent with the expectation that sequences with strongly biased amino acid compositions would dominate in the tails of the entropy map and likely be disordered (collagens being a notable exception). The overlap of folded and disordered proteins in the center of entropy map reflects the hallmark marginal stability of natural proteins where small changes in sequence can significantly alter the conformational energy landscape [31–34].

Elastin and resilin both map to the high ΔS_bb_ region of the entropy map, due to a high fractional glycine content for both proteins (**Fig. 2B, Fig. S1**). The ΔS_bb_ of elastin was less than resilin, consistent with the rubber-like recoil mechanism proposed for resilin. However, elastin also showed a lower ΔS_sol_ than resilin, counter to the hypothesis that elastin hydrophobic recoil mechanism would be reflected in a higher estimated change solvent entropy upon extension. Examination of the amino acid compositions indicates other sequence features that likely account for this discrepancy (**Fig. S1**). Alanine-rich tracts of elastin are involved in scaffolding lysines that form intermolecular crosslinks, and do not directly contribute to elastic recoil. This reduces the overall hydrophobicity of the sequence. Resilin forms materials through di-tyrosine crosslinks, and the elevated fraction of tyrosine throughout the sequence raises solvent entropy. Resilin also has a higher fraction of charged residues, which also have large ΔS_sol_ magnitudes. These charged groups may not be buried in the relaxed state and thus may not make a significant entropic contribution to recoil. Charged and polar residues can be buried in folded proteins [35–37], depending on the structural context and do contribute to folding energetics, but it is unknown whether they play an entropic role in the thermodynamics of extension for elastic protein materials. Hydration of polar residues is also a significant component of protein folding [16, 25], but their role in relaxed-extended transition is unstudied.

To determine whether the relative positions of elastin and resilin in the entropy map resulted from averaging over compositionally distinct sequence segments, we calculated backbone and solvent entropy scores for individual elastin exons and for the repetitive domains of resilin exons 1 and 3 (**Fig. 3**). Exons can represent evolutionarily exchangeable or selectable regions of sequence [38, 39], and in elastin the exon structure is well-aligned with its alternating biochemical domain architecture [40]. The higher whole-protein solvent entropy of resilin was not driven by a single atypical segment. Rather, most resilin repeats had higher solvent entropy scores than the elastin exons, indicating that the relative ordering is a distributed feature of their sequence compositions and of the residue-based solvation scale.

**Figure 3.**
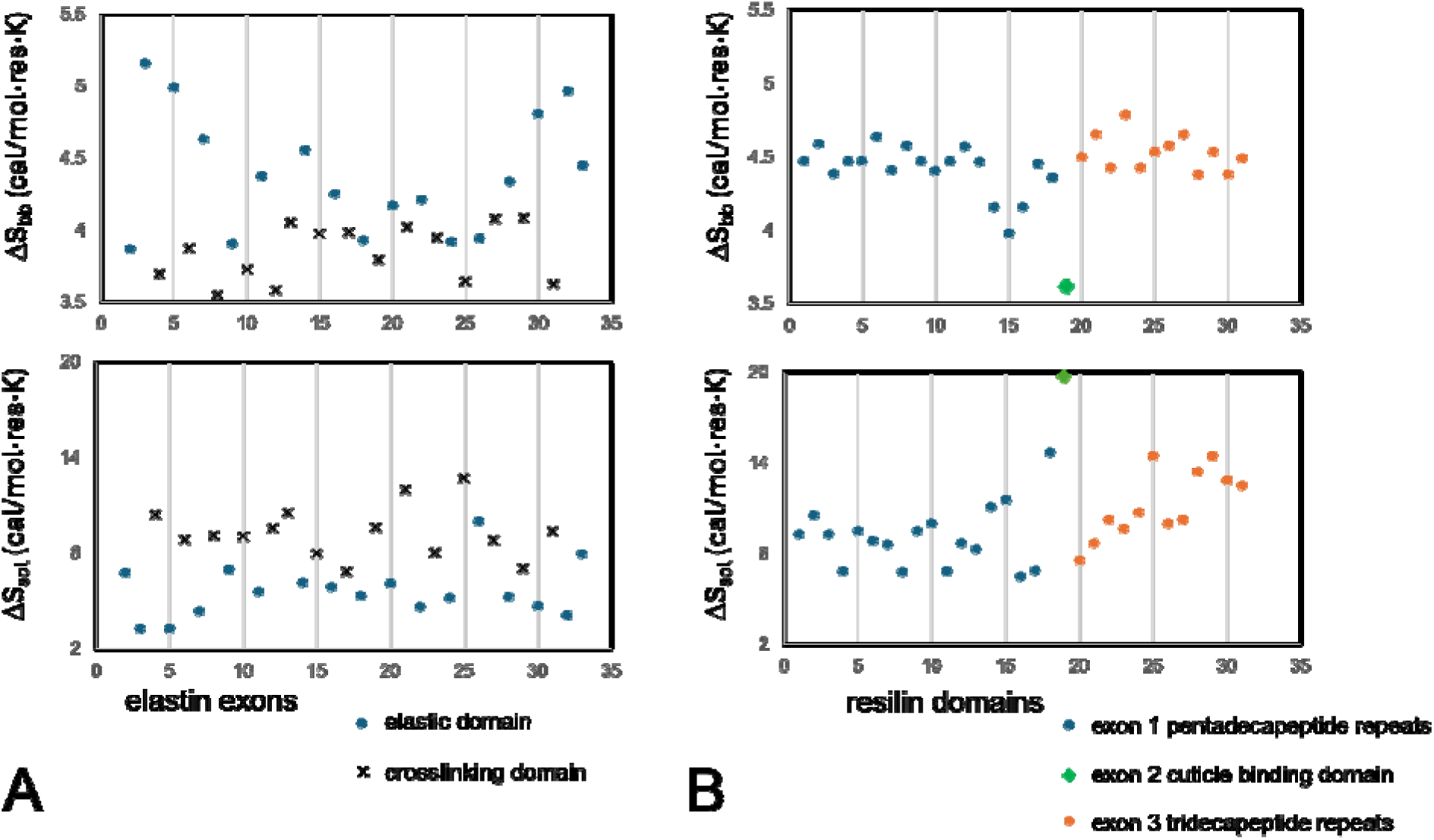
Backbone and solvent entropies for **(A)** exons in the canonical elastin isoform (UniProt P15502.3) numbered sequentially, and **(B)** *Drosophila* resilin repeat domains in exons 1 and 3 (UniProt Q9V7U0).

The two proteins also differed in the spatial organization of their entropy profiles. Resilin repeats were comparatively uniform in both backbone and solvent entropy, consistent with a mechanically repetitive material. In contrast, elastin displayed a non-uniform profile, with backbone entropy generally higher near the termini and lower across the central region, together with positional variation in solvent entropy. This organization suggests that elastin may not be mechanically homogeneous along its length, with distinct regions potentially contributing unequally to elasticity and recoil. While elastin exhibits approximately Hookean behavior over its physiological extension range, regional differences in entropy may influence the microscopic origins of that elasticity. This pattern raises the possibility that the spatial distribution of entropy, in addition to its whole-protein average, contributes to elastin mechanics. It also suggests that depending on the region of elastin under consideration, the relative contributions of hydrophobic and rubber-like recoil mechanisms may vary.

### Tissue Profiles and Putative Elastic Function

Elastin and resilin are proximal in the entropic map, suggesting that this region may include other proteins with related compositional and thermodynamic features consistent with elastic function. There are just a few proteins for which elastic function has been established, and even fewer where the thermodynamic mechanism of recoil has been studied. As a proxy for function, we investigate which proteins are specifically transcribed in mechanical tissues, and where they localize on the entropy map. Transcriptional profiles across diverse tissue types from Human Protein Atlas [41] are used to highlight proteins as ‘enhanced’ or ‘enriched’ based on tissue-specific criteria: proteins with elevated transcription levels for a particular tissue four-fold higher than the mean across all tissues are defined as tissue enhanced, and those where levels are four-fold higher than any other tissue are defined as tissue enriched. We represent these classifications as two tiers of specificity with enriched proteins more stringently tissue-specific than enhanced proteins.

In skin tissue, enriched proteins were highly overrepresented in the lower region (ΔS_sol_ < 12 cal/mol·K) of the entropy map including the high-glycine and high-proline tails (**Fig. 4A**). This is in contrast to nearly all other tissues represented in the Atlas, where both enhanced and enriched proteins are clustered near the mean of both backbone and solvent entropy (**Fig. 4B, Fig. S2).** Skin enriched proteins with ΔS_sol_ < 12 cal/mol·K cluster into two groups based on ΔSbb (**Fig. 4A**). Group 1, where ΔS_bb_ < 3.5 cal/mol·K, is less glycine-enriched with most of the proteins annotated as keratin-associated proteins (KRTAPs) (**Table S2**). In particular the KRTAP-4 family, which is involved in the hair cycle [36, 42, 43] is well represented. The KRTAP-4 family are expressed primarily in the hair cortex, where they form part of the matrix surrounding keratin intermediate filaments. They are classified as high-sulfur, due to a high sequence fraction of cysteine residues, often exceeding 30%. This is thought to enable extensive disulfide bond formation with hair keratins and other KRTAPs, producing a highly crosslinked and mechanically resilient hair fiber.

**Figure 4.**
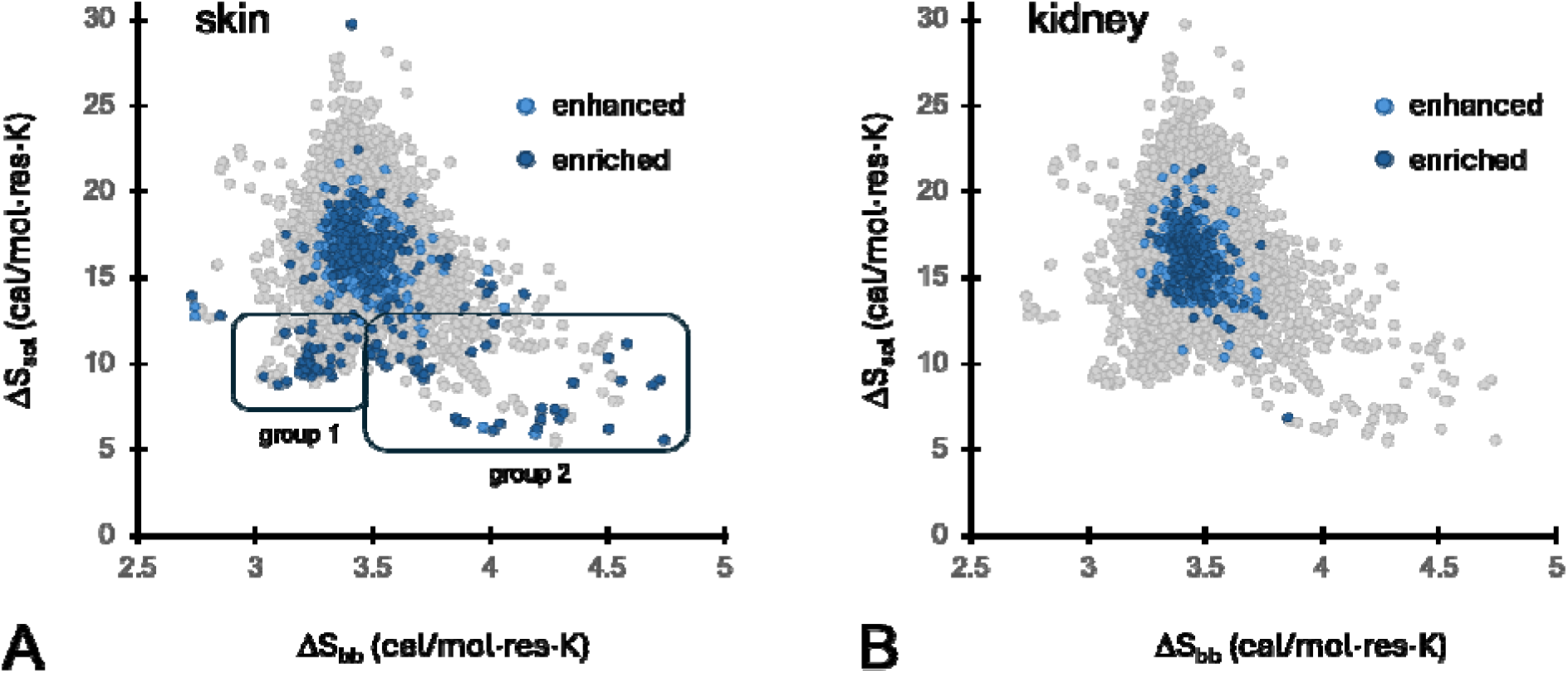
Entropy map of human proteins that are transcriptionally enhanced, four-fold over the mean, or enriched, four-fold over any other tissue in the **(A)** skin and **(B)** kidney. Maps for other tissues in the Human Protein Atlas are in **Fig. S2**.

Group 2 of skin enriched proteins, where ΔS_bb_ > 3.5 cal/mol·K, is also dominated by KRTAPs (**Fig. 4A**, **Table S2**), but instead of hair cortex mechanics, Group 2 KRTAPs are from families involved in keratinization, the mechanical maturation of the epidermis [44]. Complementing the KRTAPs are a number of late cornified envelope proteins, which contribute to cornification, the final stage of keratinization where the outer dead skin forms a tough, waterproof layer of cells filled with keratin [45]. In this group is loricrin, the major protein component of the cornified envelope [46]. Together, these proteins modulate the elastic properties of the epidermis by forming a crosslinked matrix across the skin surface [47]. Only one putative non-biomechanical protein is found in the second group, metallothionein-4, involved in zinc homeostasis in epithelial tissues [48]. Given a proposed ancient origin of KRTAPs from metallothioneins [43], it is not surprising that they share amino acid composition (high glycine, high cysteine) and map to the same region of backbone and solvent entropy.

### Neighborhoods in the Entropy Map

The criteria for enhanced and enriched protein transcript levels in tissues is stringent, and excludes proteins such as elastin which play elastic roles across multiple tissues. Elastin is a major component of the skin, forming an elastic network along with collagen [49]. We examined the neighborhood elastin in the full entropy map to see if candidates with elastic function could be identified beyond those in the skin enhanced and enriched sets. Elastin falls in the same region of the entropy map as skin Group 2 (**Fig. 2B,4A**). Its nearest neighbors are predominantly KRTAPs (**Table S3**), which are all characterized by a high fraction of cysteine (**Fig. S3**). Also highly represented are Small Cysteine and Glycine Repeat-Containing Proteins, which eponymously describe their sequence composition. Mucin-19 is found in mucus and is heavily glycosylated, altering the physiochemical properties of the residues and contributing to its viscous character. It also has a high cysteine content that allow it to form supramolecular elastic materials [50]. Other mucins were found in tissues such as the salivary gland (**Fig. S2**), although these map to Group 1, due to a high proline-content. The elastin neighborhood is enriched for disordered proteins with putative elastic function in skin, cartilage and mucus. However, given the high ΔS_bb_ and low ΔS_sol_ for this group, and the low fraction of hydrophobic residues (**Fig. S3**), it is most likely that they respond to extension through a rubber-like mechanism, rather than the hydrophobic recoil of elastin. For KRTAPs and other proteins comprising the dry environment of the epidermis, this would allow them to maintain elastic function in the absence of extensive hydration.

Resilin, while proximal to elastin, has a distinct neighborhood in the entropy map (**Fig. 2B**). Most neighbors again are KRTAPs, but differ from those adjacent to elastin. The resilin-like KRTAPs belong to high-tyrosine families. The high fraction of tyrosine in the sequence largely accounts for the greater ΔS_sol_. Although it has not been demonstrated, it is plausible that a high tyrosine content of these KRTAPs facilitates crosslinking and formation of elastic materials, much like that of resilin. Exposure of the outer layer of skin to UV-light may promote di-tyrosine crosslinking [51] during keratinization; alternatively there may be oxidases that catalyze tyrosine crosslinks, similar to the sulfhydryl oxidases that promote cysteine crosslinking in skin [52]. Specialized collagens in this neighborhood both play interfacial roles as anchors between extracellular matrix and collagen fibers in the skin (Type VII) and cartilage (Type IX) [53]. Hornerin and dermokine, are both involved in mechanical maturation of the epidermis during keratinization and cornification [54–56]. Hornerin also has a high sequence fraction of tyrosine, again perhaps playing a role in formation of crosslinked elastic networks. Assuming the high-tyrosine content of resilin neighbors is due to crosslinking, we would expect elastic function for these proteins to be driven by a rubber-like recoil mechanism.

### Entropy Map of Disordered Domains

Using the whole protein sequence to calculate ΔS_bb_ and ΔS_sol_ likely misrepresents a significant subset of the proteome that includes both ordered and intrinsically disordered regions. The proposed thermodynamic framework is predicated on a disordered ensemble in the relaxed state, and we sought to exclude folded domains from our analysis. Using the same AlphaFold DB human proteome structural dataset described earlier, we identified a set of disordered domains with a minimum length of 50 amino acids, overall pLDDT score < 0.7, containing six or fewer consecutive non-coil positions. These criteria resulted in 20,365 domains, similar in scope to standard disorder databases [57], with a mean ΔS_bb_ of 3.4 cal/mol·K and ΔS_sol_ of 16.0 cal/mol·K, both only marginally smaller than the full protein dataset (mean ΔS_bb_ = 3.5 cal/mol·K, mean ΔS_sol_ = 18.0 cal/mol·K). While a broad range of entropy space is covered, it amounts to about a third of accessible entropies, as bounded by proline, glycine and glutamic acid (**Fig. 5A**).

**Figure 5.**
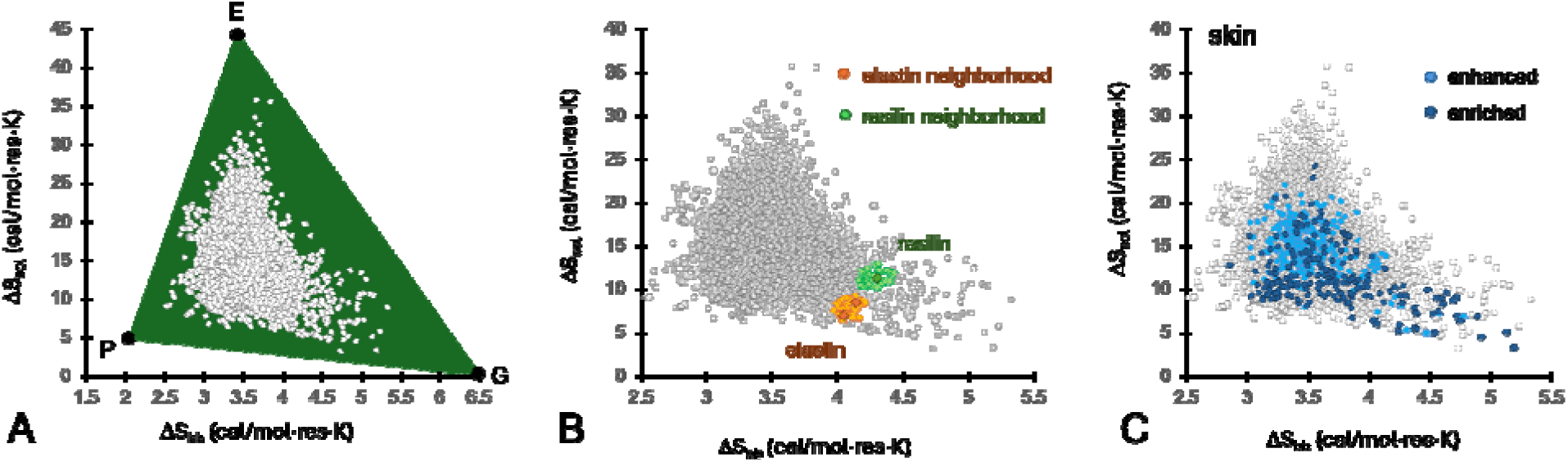
Entropy map of disordered domains of the human proteome **(A)** covers much of accessible entropy space bounded by Pro, Gly and Glu. **(B)** Euclidean nearest neighbors of elastin (N-and C-terminal domains) and resilin, and **(C)** the skin enhanced and enriched transcriptomes.

The disordered domains nearest to elastin include several proteins with established structural and biomechanical roles (**Fig. 5B,C**, **Table S5**). Most prominent are the terminal head and tail domains of epithelial type II keratins, including both termini of keratins 6A, 6B, and 6C, as well as terminal domains from keratins 4, 19, 25, 26, 75, and oral keratin 2. Keratins consist of a structured coiled-coil rod flanked by disordered terminal domains, and the recovery of these termini suggests that the entropy map identifies mechanically specialized regions that are masked by whole-protein averages. The N-terminal domains of the type II keratins are particularly relevant because a conserved lysine in the disordered head domain of keratins mediates transglutaminase-dependent attachment of keratin intermediate filaments to the cornified envelope [58]. These terminal domains are incorporated into a larger load-bearing structure through covalent crosslinking, suggesting a role for these disordered domains in supramolecular mechanics.

While the keratin domains are found at N-and/or C-termini, other elastin-neighboring domains occur internally within large extracellular matrix proteins, where they may function as compliant spacers or assembly elements between structured domains. Fibrillin-2 contains three such regions within its N-terminal portion. Fibrillin microfibrils act as scaffolds for elastin deposition and contribute limited elasticity even in elastin-free tissues [59], while amino-terminal domains of fibrillin-2 interact directly with elastin during elastic-fiber assembly [60, 61]. The identification of multiple fibrillin-2 regions near elastin is consistent with a role in accommodation, domain spacing, and load transfer within the microfibril, although elastic recoil by these individual domains has not been established. A disordered region of collagen IX is also present in the elastin neighborhood, whereas before Type IX collagen was clustered with resilin based on its whole protein sequence. Collagen IX is covalently associated with type II collagen fibrils in cartilage and functions at the interface between fibrils and the surrounding matrix, making its non-collagenous regions plausible mechanical connectors [62]. Multiple internal regions of mucin-19 were identified, and these may contribute to the extended, hydrated polymer architecture of mucus [50], although extensive glycosylation of mature mucins is expected to alter their conformational and solvation properties beyond those predicted from the unmodified amino-acid sequence.

The elastin neighborhood also includes the N-terminal disordered domains of phenylalanine/glycine-rich FG-repeat nucleoporins NUP98 and NUP54. FG-repeat nucleoporins are not conclusively established as direct mechanosensors; however, mechanical deformation of the nuclear pore complex may alter FG-Nup organization, thereby modulating transport in and out of the nucleus [63–66]. The FG-Nup proteins suggest a distinct form of disorder-driven mechanics that does not require permanent crosslinking [67, 68]. The presence of keratins, extracellular matrix connectors, mucins, and nucleoporins in the same neighborhood suggests that proximity to elastin on the entropy map may identify several architectural classes of mechanically relevant disorder: covalently anchored terminal domains, internal spacers between structured domains, and tethered polymer domains whose entropic fluctuations regulate molecular-scale organization.

The elastin disordered-domain neighborhood is enriched in domains embedded within constitutive structural assemblies where an elastic, mechanical context is supported by covalent crosslinking, stable domain attachments, or fixed tethering. In contrast, the resilin neighborhood contains a prominent group of RNA-binding and RNA-processing proteins, including FUS, EWSR1, SFPQ, hnRNP D0, and DDX3 helicases. These proteins are associated with dynamic ribonucleoprotein assemblies in which multivalent protein-protein and protein-RNA contacts act as reversible network junctions in phase separated condensates [69–71]. Their placement near resilin may therefore reflect the sequence requirements of a flexible, transiently crosslinked polymer network rather than an elastic recoil function. Notably, condensation is a critical step in the maturation of both elastin [72] and resilin [13] into a mature elastic material. Hornerin and collagen IX provide notable structural candidates within this neighborhood, as both are incorporated into mechanically constrained extracellular assemblies. Together, the two neighborhoods suggest that similar backbone and solvation entropy profiles can support distinct forms of mechanical behavior: persistent recoil in covalently integrated networks, and extension-regulated viscoelasticity in condensate phases.

Titin’s elastic recoil mechanism is not simply described by the same thermodynamic framework as elastin and resilin. Its elastic PEVK regions occupy a distinct region in the upper left of the entropy map correlated with charge-rich sequences and separate from the Gly-rich neighborhoods of elastin and resilin (**Fig. S4**). Because of titin’s exceptional length, AlphaFold DB partitions it into multiple entries corresponding to 1400 amino acid long fragments. In the whole-protein map, entries dominated by titin’s folded, immunoglobulin-like domains cluster near the center of the map, whereas entries containing substantial PEVK sequence extend toward low ΔS_bb_, high ΔS_sol_ region (**Fig. S4A**). Restricting the analysis to disordered domains removes folded Ig domains and reveals disordered titin regions distinct from the canonical PEVK cluster (**Fig. S5B**). Titin contains multiple putative classes of disordered elastic sequences. The peripheral position of PEVK is driven by its enrichment in charged residues and proline, which produces high calculated solvent entropy but relatively low backbone entropy. PEVK recoil is consequently not expected to be dominated by the Gly-dependent conformational entropy characteristic of resilin or the hydrophobic hydration mechanism of elastin. Instead, its mechanical response is likely to depend strongly on electrostatic interactions, charge patterning, ionic conditions, and polyproline-rich chain structure [73, 74], contributions that are represented only indirectly by the present two-dimensional entropy framework. Few other proteins are found in this region, but those that are, such as nebulin, share the same charge-driven elastic recoil mechanism [75].

### Comparable Models of Elastic Function

As an independent sequence-based measure of elastomeric potential, we projected the human proteome onto the glycine/proline compositional framework developed by Rauscher and colleagues (**Fig. 6**) [27]. This framework was originally proposed to distinguish elastomeric from amyloid-forming protein assemblies. Above a threshold in glycine and proline content, backbone hydration and conformational disorder are retained upon self-association, whereas sequences below this region are more permissive to intermolecular hydrogen bonding and ordered aggregation. As Rauscher et. al. showed, both elastin and resilin fall within the proposed elastomeric region, consistent with their ability to form hydrated, disordered materials despite their different recoil mechanisms.

**Figure 6.**
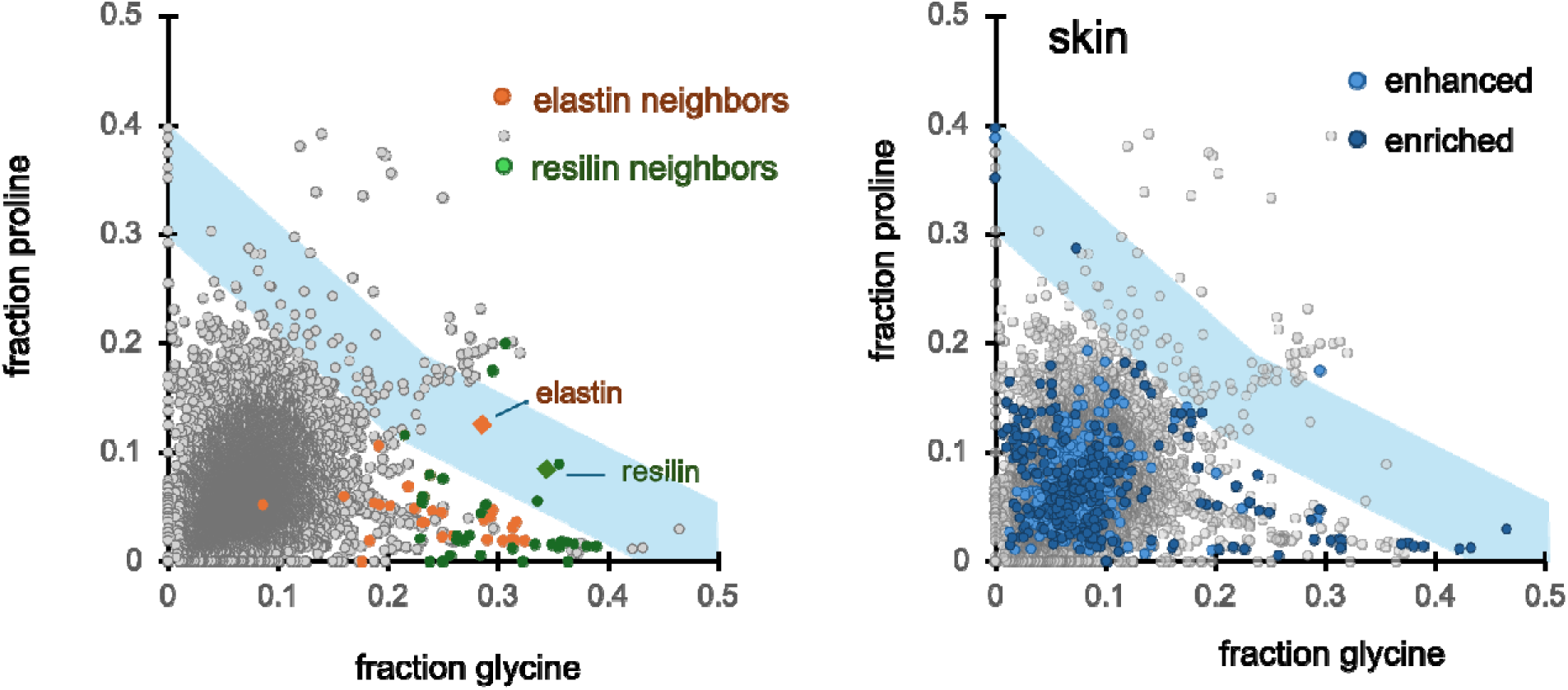
Fractional glycine and proline content was calculated for disordered domains in the human proteome. The blue-shaded region corresponds to a compositional regime associated with elastomeric rather than amyloid-like self-assembly proposed by Rauscher et al. [27]. **(A)** Disordered domains neighboring elastin and resilin on the entropy map. Elastin and resilin lie within the proposed elastomeric region, whereas most neighboring domains have lower proline content. **(B)** Disordered domains from proteins transcriptionally enhanced or enriched in skin.

In contrast, most proteins neighboring elastin and resilin on the entropy map fall outside the proposed Gly/Pro region (**Fig. 6A**). These sequences are frequently glycine-rich but contain less proline than the canonical elastomers. Their proximity in entropy space therefore does not coincide with elastomeric self-assembly based on the Gly/Pro framework. Many skin-associated domains are glycine-rich but fall below the proline content associated with autonomous elastomeric self-organization (**Fig 6B**). The Gly/Pro framework evaluates retention of hydration and conformational disorder upon assembly and is complementary to the backbone and solvent entropy coordinates used in this study. The incomplete correspondence between these representations suggests that amino acid fractions alone do not specify material behavior [67]. Proline spacing, rather than proline abundance alone, influences elastin-like self-assembly, with proline-poor intervals permitting greater local structure and altered aggregation behavior. Likewise, charge patterning, aromatic sticker spacing, crosslinking sites, and domain position can determine whether a disordered sequence forms an elastic network, a viscoelastic condensate, or an ordered aggregate.

While overall amino acid composition captures important thermodynamic features of elastic proteins, functional behavior is also influenced by sequence grammar, namely the ordering and patterning of residues along the chain [76]. Ruff and colleagues analyzed the human disordered proteome for sequence-pattern features and identified grammar classes associated with different molecular behaviors [4]. In their framework, cluster 22 is enriched in well-mixed glycine-and proline-rich sequences and is proposed to represent elastomeric proteins, consistent with the enrichment of known elastic proteins in this compositional regime [27] (**Fig. S5**). Cluster 22 maps to the high ΔS_bb_, low ΔS_sol_ region of the entropy map that contains elastin and resilin-like sequences and many candidate disorder-based elastic proteins. This cluster mostly comprises collagens. It does not include KRTAPs or any of the proteins involved in the hair cycle or keratinization. Elastin itself is not in cluster 22. Instead, a disordered fragment of elastin exon 26 maps to a cluster characterized by blocks of polar residues (**Fig. S5**). This cluster is more centrally located on entropy map than elastin suggesting that elastin-like behavior emerges from sequence architectures distinct from the canonical Gly/Pro-rich elastomer grammar.

### Sequence Grammar of Elastic Function in Protein Language Models

Protein language models (PLMs) are trained on hundreds of millions of protein sequences and learn statistical representations that capture evolutionary, structural, and biophysical relationships directly from sequence [77, 78]. Rather than requiring explicit specification of sequence motifs or residue patterning rules, PLM embeddings provide a high-dimensional representation in which proteins with related functional constraints often cluster together despite limited sequence similarity [79, 80]. We reasoned that if elastic recoil mechanisms are encoded by subtle combinations of composition, sequence grammar, and evolutionary context, then these features may be reflected in PLM latent space. This approach provides an opportunity to identify elastic protein classes beyond those recognized by current grammar-based models and to determine whether proteins sharing common thermodynamic recoil mechanisms occupy distinct regions of sequence embedding space.

To determine whether elastic proteins occupy distinct regions of protein sequence space beyond those identified by explicit grammar rules, we embedded all disordered domains using the ESM2 protein language model with 650 million parameters [77]. ESM2 representations are learned from large-scale self-supervised training on protein sequence databases and capture information related to evolutionary constraint, structural organization, and biochemical function without requiring predefined rules or alignments. Rather than relying on a single representation layer, we extracted mean pooled sequence embeddings from layers 24, 30, and 33 and concatenated them into a composite feature vector. Different layers of transformer language models encode different levels of biological information, with intermediate layers often capturing local biochemical and structural features while deeper layers increasingly reflect higher-order functional and evolutionary relationships [81]. By integrating information across multiple layers, we sought to capture a broader representation of sequence grammar than is available from any single embedding. The resulting embedding space was analyzed using complementary graph-based approaches. A k-nearest-neighbor (kNN) graph was used to define local relationships among proteins based on cosine similarity [82], preserving neighborhoods that reflect fine-scale similarities in sequence organization and biophysical properties. Leiden community detection was then applied to identify densely connected groups of proteins within the graph, providing a hierarchical view of larger functional and evolutionary neighborhoods [83]. Together, these approaches allow both local and global features of manifold topology to be examined, revealing whether proteins with shared thermodynamic recoil mechanisms cluster into common latent-space communities despite divergent amino acid composition or sequence grammar.

Projection of the ESM2 embeddings into two dimensions using UMAP [84] revealed a structured latent space in which most Leiden communities formed coherent neighborhoods, although several communities exhibited partial overlap or separation into distinct islands (**Fig. 7**). Community sizes were heterogeneous, indicating that the sequence features learned by the protein language model are unevenly distributed across the disordered proteome. Communities enriched in highly biased sequences were generally more compact and isolated, whereas larger communities occupied central regions of the manifold and showed greater overlap with neighboring groups. This pattern is consistent with the expectation that extreme compositional and grammatical features, such as the glycine-rich repeats characteristic of resilin-like proteins, generate stronger sequence signatures than more heterogeneous elastic proteins. A total of sixteen Leiden communities were identified, with elastin (exon 26), resilin (residues 39-108), and the PEVK (residues 9787-9906) region of titin each partitioning into distinct communities. The separation of these canonical elastic systems suggests that the language model captures sequence features associated with different modes of elastic recoil rather than grouping all elastic proteins into a single class. At the same time, the proximity of some communities within the manifold suggests the presence of intermediate sequence grammars that may connect established elastic protein families to previously unrecognized mechanically functional proteins. Together, these results indicate that protein language models encode aspects of elastic protein sequence organization beyond simple amino acid composition and can distinguish among multiple thermodynamic and biomechanical solutions to elastic function.

**Figure 7.**
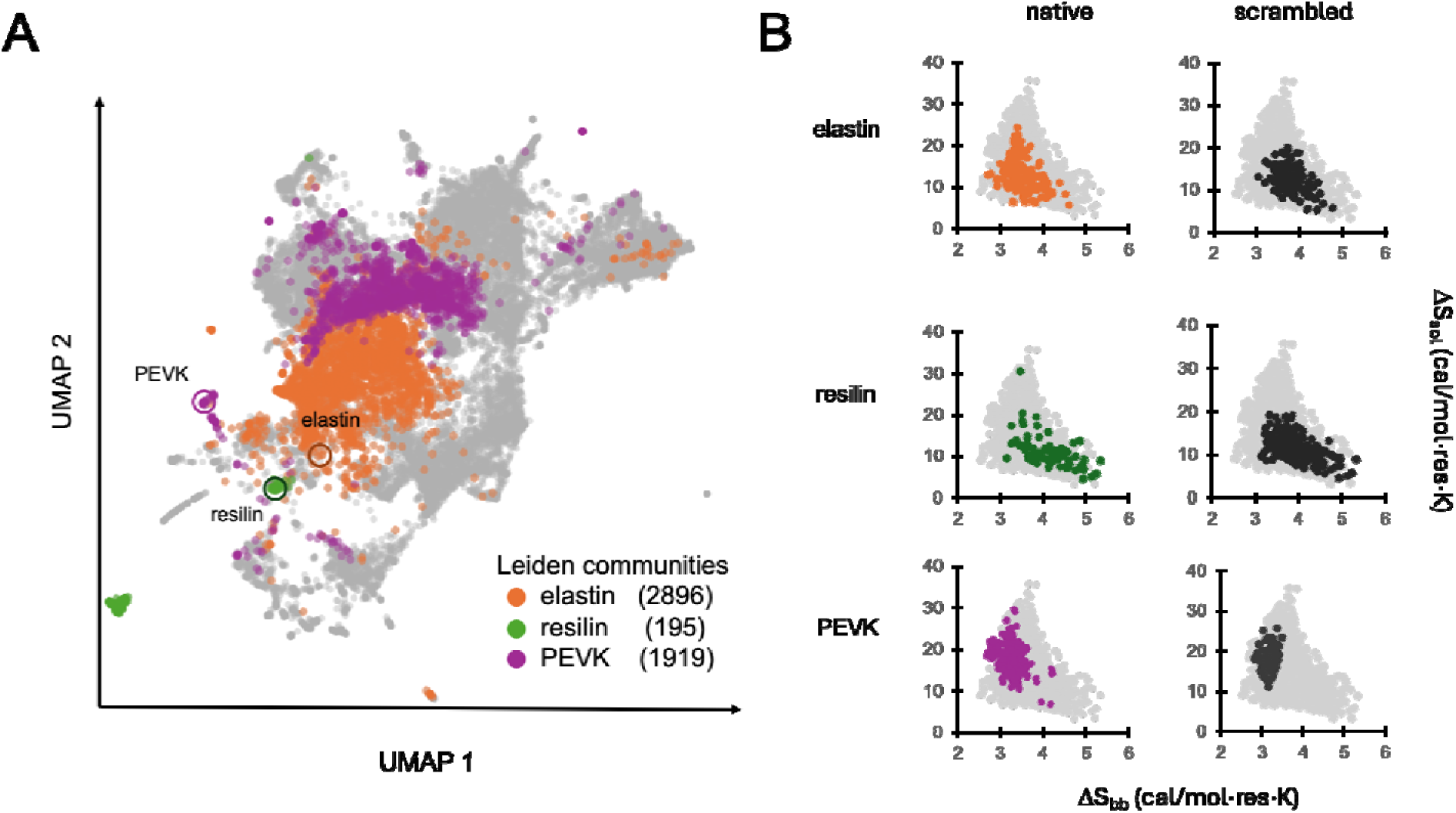
Protein language model embedding space of disordered domains. Disordered domains from the human proteome were embedded using ESM2-650M representations generated from multiple layers. **(A)** UMAP projection of the concatenated embeddings reveals coherent Leiden community structure. Canonical elastic proteins occupy distinct communities. **(B)** kNN neighborhoods from the full manifold map to distinct regions of the backbone-solvent entropy landscape.

Although elastin, resilin, and the titin PEVK region each occupied distinct Leiden communities, there was not a strict correspondence between Leiden community assignment and nearest-neighbor relationships in embedding space. In several cases, proteins identified as nearest neighbors by the kNN analysis crossed Leiden community boundaries, indicating that local and regional features of the manifold capture different aspects of sequence organization. The kNN neighborhoods reflect fine-scale similarity in the embedding space and are sensitive to local sequence grammars, whereas Leiden communities identify broader regions of the manifold defined by dense connectivity and shared higher-order sequence features. Despite these differences, the nearest-neighbor sets of elastin-, resilin-, and PEVK-like domains showed clear organization when projected onto the entropy map (Fig. 7). Neighboring sequences frequently occupied thermodynamic regions consistent with the canonical protein around which they clustered, with elastin neighbors enriched in low ΔS_sol_, high ΔS_bb_ disorder-rich proteins, resilin neighbors concentrated in a distinct glycine-rich region of entropy space, and PEVK-associated sequences extending toward the charge-rich regime of elevated ΔS_sol_. The agreement between thermodynamic proximity and local relationships in PLM embedding space suggests that the model captures features related to the physical basis of elastic recoil. At the same time, the imperfect overlap between kNN neighborhoods and Leiden communities indicates that elastic proteins are embedded within broader sequence families whose shared grammatical features may extend beyond the specific thermodynamic mechanisms highlighted by the entropy framework.

The amino acid compositions of kNN networks for elastin and resilin are consistent with their putative recoil mechanisms (**Fig. S6**). The resilin neighborhood is strongly enriched in glycine, tyrosine, glutamine, asparagine, and arginine, consistent with the prevalence of low-complexity RNA-binding proteins, condensate-associated scaffolds, and glycine-rich repetitive sequences. In contrast, the elastin neighborhood is enriched in proline, alanine, valine, and other hydrophobic residues characteristic of classical elastomeric and extracellular matrix proteins. These compositional differences mirror the distinct thermodynamic and sequence-grammar signatures of the two protein classes, with resilin-associated networks favoring highly disordered, interaction-rich polymers and elastin-associated networks favoring hydrophobic, entropically elastic architectures.

The PLM neighborhood of resilin was enriched in structural proteins associated with mechanically resilient tissues: LDB3, COL17A1, loricrin, and multiple keratins, all of which contribute to mechanically robust filamentous or extracellular matrix architectures (**Table S7**). In parallel, the neighborhood contained numerous low-complexity RNA-binding proteins, including FUS, DDX3 helicases, RBM14, and several hnRNP family members. These proteins are central components of dynamic ribonucleoprotein assemblies whose organization depends on intrinsically disordered regions and reversible multivalent interactions [68]. The coexistence of cytoskeletal, extracellular matrix, and condensate-associated proteins within the resilin neighborhood suggests that the PLM is detecting sequence features associated with deformable polymer networks across multiple biological scales, ranging from tissue mechanics to intracellular biomolecular assemblies.

The PLM-derived kNN neighborhood of elastin reveals a mixture of known matrix proteins, mechanically associated scaffolds, and regulatory proteins containing extensive intrinsically disordered regions (**Table S8**). The most notable neighbors are fibrillin-1 (FBN1) and fibrillin-2 (FBN2), both of which are established contributors to elastic fiber architecture and are annotated as extracellular matrix constituents conferring elasticity, providing an important validation that the PLM embedding space recovers proteins involved in biological elasticity [59–61]. Additional extracellular matrix proteins, including collagen VIII (COL8A1), as well as cytoskeletal and adhesion-associated proteins such as IFFO1, WASF1, and CNTNAP1, suggest enrichment for proteins that function within mechanically active cellular or extracellular networks [85–88]. The neighborhood of elastin contains not only established elastic-fiber proteins but also a diverse collection of scaffold, RNA-binding, and transcriptional regulatory proteins containing extensive intrinsically disordered regions. Although these proteins are not generally classified as elastic, several belong to classes in which disordered segments have been proposed to function as entropic springs, flexible tethers, or molecular spacers. For example, large regulatory scaffolds such as SPEN and BAG6 contain long disordered linkers separating multiple interaction domains, architectures that could permit reversible extension and contraction while maintaining connectivity between distant binding partners in processes like chromatin regulation or protein quality control [89–91]. Likewise, RNA-binding proteins (NOVA1, MBNL2, PTBP3, and UPF1) that participate in dynamic ribonucleoprotein assemblies may mediate transient interactions while preserving molecular mobility. More generally, intrinsically disordered regions may function as mechanically active elements that can generate entropic restoring forces, regulate intermolecular spacing, and buffer structural fluctuations within large macromolecular assemblies. The recovery of proteins with related architectural features in the elastin neighborhood therefore suggests that sequence determinants underlying elastic behavior may be reused across diverse cellular contexts. Rather than being restricted to specialized extracellular elastomers, elastic-like sequence grammars may contribute more broadly to molecular flexibility, mesoscale organization, and adaptive mechanics [92].

To assess the extent to which PLM-derived neighborhoods depend on sequence grammar rather than amino acid composition alone, we generated ten independently scrambled variants of the elastin exon 26 query and ten scrambled variants of the analyzed resilin domain while preserving overall amino acid composition. The entropy maps of the scrambled sequence kNNs are similar to the native sequence, indicating that composition is a strong signal for similarity in these sequences. However, among the protein neighbors recovered across the ten scrambled elastin searches, only a single protein, zinc finger protein 865, was also recovered by the native elastin query (**Table S8**), indicating that the elastin neighborhood is highly sensitive to residue ordering. In contrast, scrambled resilin sequences recovered several proteins from the native resilin neighborhood, with several proteins appearing in multiple independent scrambles (**Table S7**). The persistence of these relationships despite sequence randomization suggests that the resilin neighborhood is determined in part by compositional and low-complexity sequence features that survive scrambling. One explanation is that resilin occupies an extreme low-complexity region of sequence space dominated by glycine-, proline-, serine-and tyrosine-rich sequence statistics, making many randomized variants remain recognizably resilin-like to the language model.

## CONCLUSIONS

We explored a sequence-derived thermodynamic framework to organize intrinsically disordered proteins according to their putative mechanisms of elastic recoil. The entropy landscape separates canonical elastic proteins such as elastin and resilin, while also identifying numerous candidate proteins enriched in extracellular matrices, cytoskeletal assemblies, and dynamic molecular scaffolds. Elastic function may be substantially more widespread than currently recognized and the relative contributions of backbone and solvent entropy provide a useful framework for classifying candidate elastic proteins. At the same time, the present model has important limitations. The solvent-entropy term is estimated solely from amino acid composition and implicitly assumes extension-induced changes in accessibility, whereas the thermodynamics of real disordered proteins depends on the conformational ensemble they sample in solution. Consequently, proteins with similar compositions but different sequence grammars or local structural preferences may occupy similar positions in the entropy map despite possessing distinct physical mechanisms. Future refinements should incorporate information from molecular simulations, polymer models, experimental ensemble measurements, or learned representations of conformational heterogeneity to better estimate the solvent environment experienced by residues in the relaxed state and thereby improve mechanistic discrimination among elastic proteins.

The thermodynamic variables governing elastic recoil may also be related to the phase behavior of intrinsically disordered proteins. Elastin-like polypeptides exhibit heat-induced phase separation driven by release of hydration water [93], a process closely related to the solvent-entropy mechanism proposed here for elastin recoil. In contrast to elastin-like heat-induced transitions that are closely linked to dehydration and solvent entropy, many resilin-like proteins and biomolecular condensates undergo associative phase transitions driven by favorable intermolecular interactions, including electrostatic, aromatic, and hydrogen-bonding contacts [94]. These interaction-driven transitions are conceptually closer to cooling-induced behavior, where enthalpic stabilization of the condensed phase competes against mixing entropy. Incorporation of residue-specific enthalpic descriptors may therefore provide a means to extend the present entropy framework toward prediction of both elastic recoil and phase behavior [95].

The neighborhoods of elastin and resilin in the PLM and entropy maps highlight a conceptual challenge in defining elastic proteins as a distinct functional category. Many proteins recovered near elastin or resilin are not known to perform overt biomechanical roles, yet contain extensive intrinsically disordered regions that may act as molecular tethers, flexible linkers, condensate scaffolds, or components of mechanically resilient cellular architectures. In such systems, elastic recoil may contribute only indirectly to biological function by regulating spacing, adaptability, or the dynamics of macromolecular assemblies. This blurs the boundary between elastic function and intrinsic disorder itself, making it challenging to establish a strict classification of elastic proteins based solely on known biological function. However, this ambiguity may also provide insight into the evolutionary origins of elastic systems. Rather than emerging *de novo*, specialized elastomers may evolve through amplification, repetition, or repurposing of disordered sequence motifs that originally served regulatory, structural, or interaction-mediated functions. From this perspective, canonical elastic proteins such as elastin and resilin may represent highly optimized endpoints along a broader continuum of disordered protein behavior in which entropic elasticity is a latent property that can be co-opted and refined for mechanical function.

## CODE AVAILABILITY

All code used for entropy calculations, disordered-domain identification, protein language model embedding generation, nearest-neighbor analyses and Leiden community detection is available at: https://github.com/vikasnanda/elastic-proteome-thermodynamics

## ACKNOWLEDGEMENTS

VN was supported by a NASA Astrobiology Institute Grant 80NSSC18M0093 and a NASA Exobiology Grant 80NSSC24K1165. We thank Benjamin Schuster and Keith Mickolajczyk for feedback on the manuscript.

## SUPPLEMENTARY MATERIALS

### METHODS

#### Structural Database

Predicted protein structures were obtained from the AlphaFold Protein Structure Database (AlphaFold DB), which provides proteome-scale structural predictions generated using AlphaFold2 and indexed by UniProt accession [29, 30]. We analyzed the complete human reference proteome (downloaded from the EMBL-EBI AlphaFold DB repository (UP000005640_9606_v6). For most proteins, a single AlphaFold model is provided for the full-length sequence. However, proteins longer than approximately 2,700 amino acids are distributed as multiple overlapping structure fragments because of AlphaFold model length limitations. In AlphaFold DB, these proteins are represented as separate entries corresponding to overlapping sequence windows of up to 1,400 residues. This includes the elastic protein titin. For analyses performed at the whole-protein level, each AlphaFold DB fragment was treated as an independent structural entry. For subsequent domain-level analyses, intrinsically disordered regions were identified independently within each fragment and retained with their associated UniProt accession and residue numbering. This treatment allowed uniform processing of both standard-length proteins and very large multidomain proteins while maintaining direct traceability to the original AlphaFold DB records.

#### Elastin exons

Elastin exons are based on the canonical protein sequence, UniProt entry P15502.3. Exons are numbered sequentially. Crosslinking domains are classified based on presence of Ala/Lys or Pro/Lys motifs.

~~~
>P15502.3.1 N-terminal domain
MAGLTAAAPRPGVLLLLLSILHPSRPGG
>P15502.3.2 hydrophobic elastic domain
VPGAIPGGVPGGVFYPG
>P15502.3.3 hydrophobic elastic domain
AGLGALGGGA
>P15502.3.4 crosslinking domain
LGPGGKPLKPV
>P15502.3.5 hydrophobic elastic domain
PGGLAGAGLGAG
>P15502.3.6 crosslinking domain
LGAFPAVTFPGALVPGGVADAAAAYKAAKAG
>P15502.3.7 hydrophobic elastic domain
AGLGGVPGVGGLGVSAG
>P15502.3.8 crosslinking domain
AVVPQPGAGVKPGKVPG
>P15502.3.9 hydrophobic elastic domain
VGLPGVYPGGVLPG
>P15502.3.10 crosslinking domain
ARFPGVGVLPGVPTGAGVKPKAPG
>P15502.3.11 hydrophobic elastic domain
VGGAFAGIPG
>P15502.3.12 crosslinking domain
VGPFGGPQPGVPLGYPIKAPKLPG
>P15502.3.13 crosslinking domain
GYGLPYTTGKLPYG
>P15502.3.14 hydrophobic elastic domain
YGPGGVAGAAGKAGYPTGTG
>P15502.3.15 crosslinking domain
VGPQAAAAAAAKAAAKFG
>P15502.3.16 hydrophobic elastic domain
AGAAGVLPGVGGAGVPGVPGAIPGIGGIAGE
>P15502.3.17 crosslinking domain
VGTPAAAAAAAAAAKAAKYG
>P15502.3.18 hydrophobic elastic domain
AAAGLVPGGPGFGPGVVGVPGAGVPGVGVPGAGIPVVPGAGIP
>P15502.3.19 crosslinking domain
GAAVPGVVSPEAAAKAAAKAAKYG
>P15502.3.20 hydrophobic elastic domain
ARPGVGVGGIPTYGVGAGGFPGFGVGVGGIPGVAGVPGVGGVPGVGGVPGVGISP
>P15502.3.21 crosslinking domain
EAQAAAAAKAAKYG
>P15502.3.22 hydrophobic elastic domain
AAGAGVLGGLVPGAPGAVPGVPGTGGVPG
>P15502.3.23 crosslinking domain
VGTPAAAAAKAAAKAAQFG
>P15502.3.24 hydrophobic elastic domain
LVPGVGVAPGVGVAPGVGVAPGVGLAPGVGVAPGVGVAPGVGVAPGIGPGGVAA
>P15502.3.25 crosslinking domain
AAKSAAKVAAKAQLR
>P15502.3.26 hydrophobic elastic domain
AAAGLGAGIPGLGVGVGVPGLGVGAGVPGLGVGAGVPGFGAGADEGVRRSLSPELREGDPSSSQ HLPSTPSSPRV
>P15502.3.27 crosslinking domain
PGALAAAKAAKYG
>P15502.3.28 hydrophobic elastic domain
AAVPGVLGGLGALGGVGIPGGVVG
>P15502.3.29 crosslinking domain
AGPAAAAAAAKAAAKAAQFG
>P15502.3.30 hydrophobic elastic domain
LVGAAGLGGLGVGGLGVPGVGGLGG
>P15502.3.31 crosslinking domain
IPPAAAAKAAKYG
>P15502.3.32 hydrophobic elastic domain
AAGLGGVLGG
>P15502.3.33 hydrophobic elastic domain
AGQFPLGG
>P15502.3.34 C-terminal domain 1
VAARPGFGLSPIFPG
>P15502.3.35 C-terminal domain 2
GACLGKACGRKRK
~~~

#### Resilin domains

Resilin domains are based on the canonical sequence, UniProt Q9V7U0. Pentadecapeptide units in exon 1 and tricadecapeptide units in exon 3 were defined by manual sequence alignments.

~~~
>Q9V7U0.1
GGRPSDSYGAPGGGN
>Q9V7U0.2
GGRPSDSYGAPGQGQGQGQGQGGY
>Q9V7U0.3
AGKPSDTYGAPGGGNGN
>Q9V7U0.4
GGRPSSSYGAPGGGN
>Q9V7U0.5
GGRPSDTYGAPGGGN
>Q9V7U0.6
GGRPSDTYGAPGGGGNGN
>Q9V7U0.7
GGRPSSSYGAPGQGQGNGN
>Q9V7U0.8
GGRSSSSYGAPGGGN
>Q9V7U0.9
GGRPSDTYGAPGGGN
>Q9V7U0.10
GGRPSDTYGAPGGGNN
>Q9V7U0.11
GGRPSSSYGAPGGGN
>Q9V7U0.12
GGRPSDTYGAPGGGNGNGS
>Q9V7U0.13
GGRPSSSYGAPGQGQGGF
>Q9V7U0.14
GGRPSDSYGAPGQ
>Q9V7U0.15
NQKPSDSYGAPGSGNGN
>Q9V7U0.16
GGRPSSSYGAPGSGP
>Q9V7U0.17
GGRPSDSYGPPASGSGAGGAG
>Q9V7U0.18
GSGPGGADYDND
>Q9V7U0.19
EPAKYEFNYQVEDAPSGLSFGHSEMRDGDFTTGQYNVLLPDGRKQIVEYEADQQGYRPQIRYEGDANDGS
>Q9V7U0.20
GPSGPGGPGGQNLGAD
>Q9V7U0.21
GYSSGRPGNGNGNGNG
>Q9V7U0.22
GYSGGRPGGQDLGPS
>Q9V7U0.23
GYSGGRPGGQDLGAG
>Q9V7U0.24
GYSNGKPGGQDLGPG
>Q9V7U0.25
GYSGGRPGGQDLGRD
>Q9V7U0.26
GYSGGRPGGQDLGAS
>Q9V7U0.27
GYSNGRPGGNGNGGSD
>Q9V7U0.28
GGRVIIGGRVIGGQDGGDQ
>Q9V7U0.29
GYSGGRPGGQDLGRD
>Q9V7U0.30
GYSSGRPGGRPGGNGQDSQDGQ
>Q9V7U0.31
GYSSGRPGQGGRNGFGPGGQNGDNDGSGYRY
~~~

#### Entropy Map Generation and Neighbor Selection

Per-residue backbone and solvent entropy scores for a protein or domain were computed by summing across sequence and normalizing for sequence length *N*.

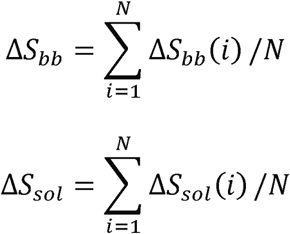

Neighbors were identified by ranking sequences according to Euclidean distance in the entropy map. Tools for generating an entropy map from sequence are included in the code repository.

#### Human Protein Atlas

Tissue-specific transcriptomes were accessed from the Human Protein Atlas [41] in July 2026. Tissue-enhanced and tissue-enriched annotations were used to evaluate whether candidate elastic proteins were preferentially associated with mechanically active tissues. Protein records were mapped to UniProt accessions and merged with the AlphaFold DB-derived protein dataset using UniProt identifiers.

#### Protein Language Model Embedding Calculations for Intrinsically Disordered Domains

Intrinsically disordered domains were represented using embeddings generated by the ESM2 protein language model (ESM2-t33-650M-UR50D) [77], a transformer-based network trained on large-scale protein sequence databases using self-supervised masked language modeling. ESM2 contains approximately 650 million parameters and has been shown to learn biologically meaningful representations that capture sequence, structural, and functional relationships directly from primary sequence. Sequences corresponding to disordered domains were derived from AlphaFold DB human proteome entries and provided as amino acid fragments with associated UniProt accession, residue boundaries, and sequence length. Protein language model embeddings were generated using the FAIR implementation of ESM2. The model was executed in inference mode without gradient calculation using either CUDA-enabled GPUs or CPU hardware depending on availability. Code is available in the github repository.

Prior to embedding generation, sequences were stripped of non-standard characters, and ambiguity codes (X, B, Z, U, O) were removed. Sequence lengths were recalculated following cleaning, and empty entries were discarded. Fragment identifiers were generated by concatenating UniProt accession and residue coordinates to maintain traceability between sequence fragments and embedding outputs. ESM2 supports a maximum sequence length of 1,022 residues and all disordered domains in the analyzed dataset were substantially shorter than this limit. For each sequence, transformer representations were extracted from layers 24, 30, and 33. These layers were selected to capture information from multiple depths within the transformer architecture, as previous analyses of protein language models have demonstrated that intermediate and deep layers encode distinct biological features, including local biochemical properties, secondary-structure information, long-range sequence dependencies, and higher-order functional relationships.

For each extracted layer, residue-level embedding vectors were obtained from the ESM2 hidden representations. Beginning-of-sequence and end-of-sequence special tokens were excluded from subsequent calculations. A single fixed-length embedding for each domain was generated by mean pooling across all residue embeddings:

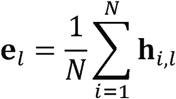

where h is the hidden-state vector for residue i at transformer layer l, and *N* is the number of amino acid residues in the fragment. This generates a 1,280-dimensional embedding vector for each sequence at each of the selected layers.

Mean-pooled embeddings were stored independently for layers 24, 30, and 33 as 32-bit floating-point matrices with dimensions *n* × 1280, where *n* corresponds to the number of analyzed disordered domains. Fragment metadata, including UniProt accession, residue coordinates, sequence length, and fragment identifier, were retained in parallel for downstream integration. Layer-specific embeddings were saved independently and additionally archived in a compressed multi-layer format to facilitate downstream manifold learning and clustering analyses.

For subsequent analyses, embeddings from layers 24, 30, and 33 were L2-normalized individually and concatenated to generate a composite representation spanning multiple levels of learned sequence abstraction. The resulting embedding space was used for dimensionality reduction, manifold reconstruction, nearest-neighbor identification, and Leiden community analysis.

#### PLM Nearest Neighbor Analysis

To identify proteins occupying similar regions of protein language model (PLM) embedding space as canonical elastic proteins, representative query sequences from elastin (ELN; UniProt P15502), Drosophila resilin (UniProt Q9V7U0), and the PEVK region of titin (TTN; UniProt Q8WZ42) were analyzed against the complete human disordered-domain embedding dataset. Query sequences were selected from experimentally characterized elastic regions and cleaned using the same preprocessing pipeline applied to the human disordered-domain dataset. Query sequences were embedded using the ESM2-t33-650M-UR50D protein language model under the same conditions used for the human disordered-domain database. Query embeddings were concatenated with the precomputed human disordered-domain embeddings to generate a common embedding matrix. This combined dataset allowed query sequences and human domains to be analyzed simultaneously within the same latent space.

Local similarity relationships within the combined embedding space were reconstructed using a k-nearest-neighbor (kNN) graph. Pairwise distances were computed using cosine distance on the normalized embedding vectors. For each sequence, the 50 nearest neighbors were identified using brute-force nearest-neighbor search. The resulting graph was represented as an undirected weighted network in which vertices corresponded to protein fragments and edges connected neighboring sequences. Edge weights were defined as *w* = 1 – *d_cos_*, where *d_cos_*is the cosine distance between two embedding vectors. Consequently, highly similar proteins received larger edge weights, while more distant neighbors contributed weaker connections. Duplicate edges were collapsed by retaining the maximum observed edge weight.

Domains were ranked according to increasing cosine distance from the query. The top 100 nearest fragments were retained for fragment-level analyses. To reduce redundancy resulting from multiple disordered domains occurring within a single protein, protein-level rankings were generated by selecting the closest fragment associated with each UniProt accession and then ranking proteins according to their minimum query distance. The top 50 unique proteins were retained as query-associated neighbors.

#### Community Detection

Broader organization of the PLM manifold was assessed using Leiden community detection on the weighted kNN graph. Communities were identified using the RBConfigurationVertexPartition implementation [83] with a resolution parameter of 1.0. The Leiden algorithm partitions the graph into densely connected communities while maximizing modularity and ensuring well-connected clusters. Community labels were subsequently assigned to all query sequences and human disordered domains.

#### Dimensionality Reduction and Visualization

For visualization, the combined embedding matrix was first reduced using principal component analysis (PCA) to a maximum of 50 components. A two-dimensional Uniform Manifold Approximation and Projection (UMAP) representation [84] was subsequently generated using cosine distance, 50 neighbors, and a minimum distance parameter of 0.1. UMAP coordinates were used solely for visualization and did not contribute to nearest-neighbor calculations. All nearest-neighbor rankings were computed in the original high-dimensional ESM2 embedding space.

## SUPPLEMENTARY FIGURES

**Figure S1.**
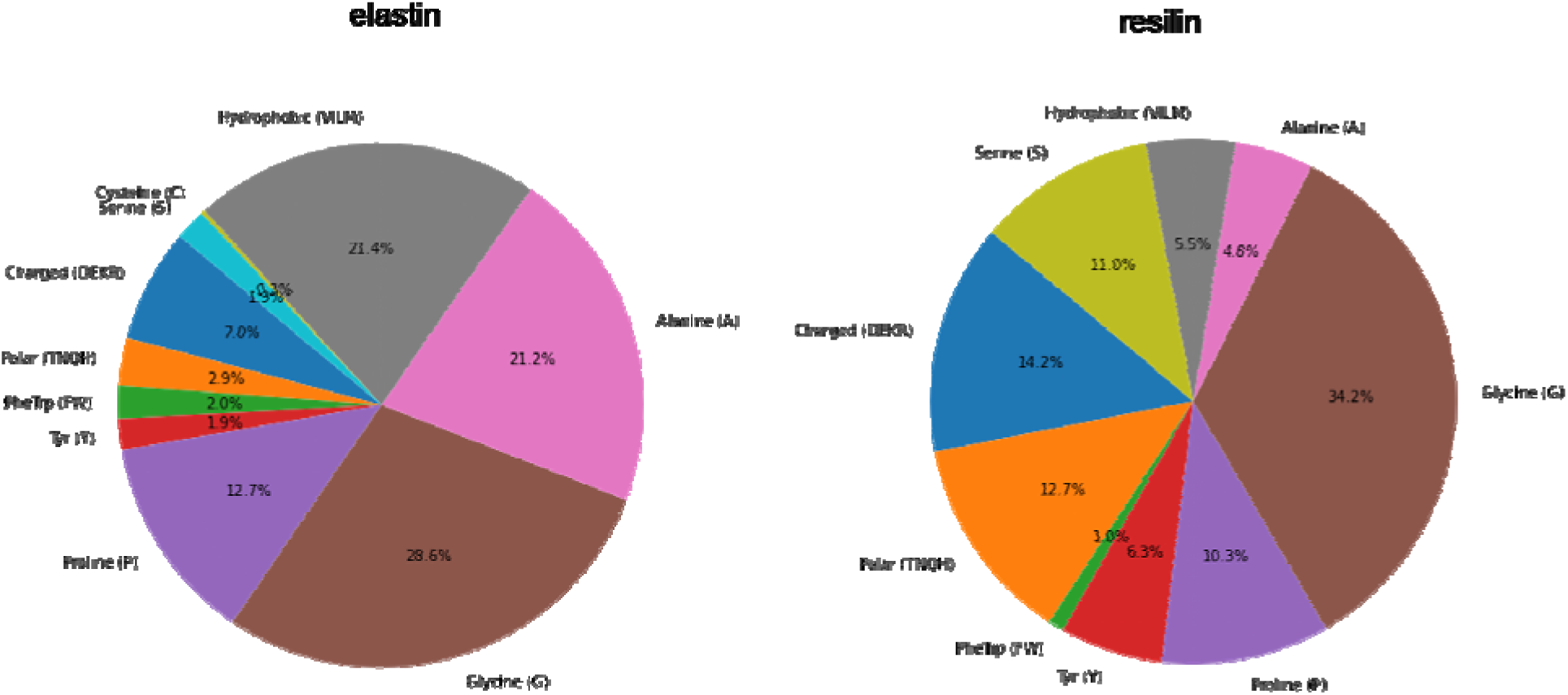
Amino acid compositions of elastin and resilin. Both elastic proteins have large fractions of proline and glycine. The key differences are in fraction polar/charged (resilin 37.9%, elastin 19.7%) versus hydrophobic (resilin 5.5%, elastin 21.4%). Resilin has a high fraction of tyrosines, involved in crosslinking. Elastin has a high fraction of alanines that scaffold crosslinking regions.

**Figure S2.**
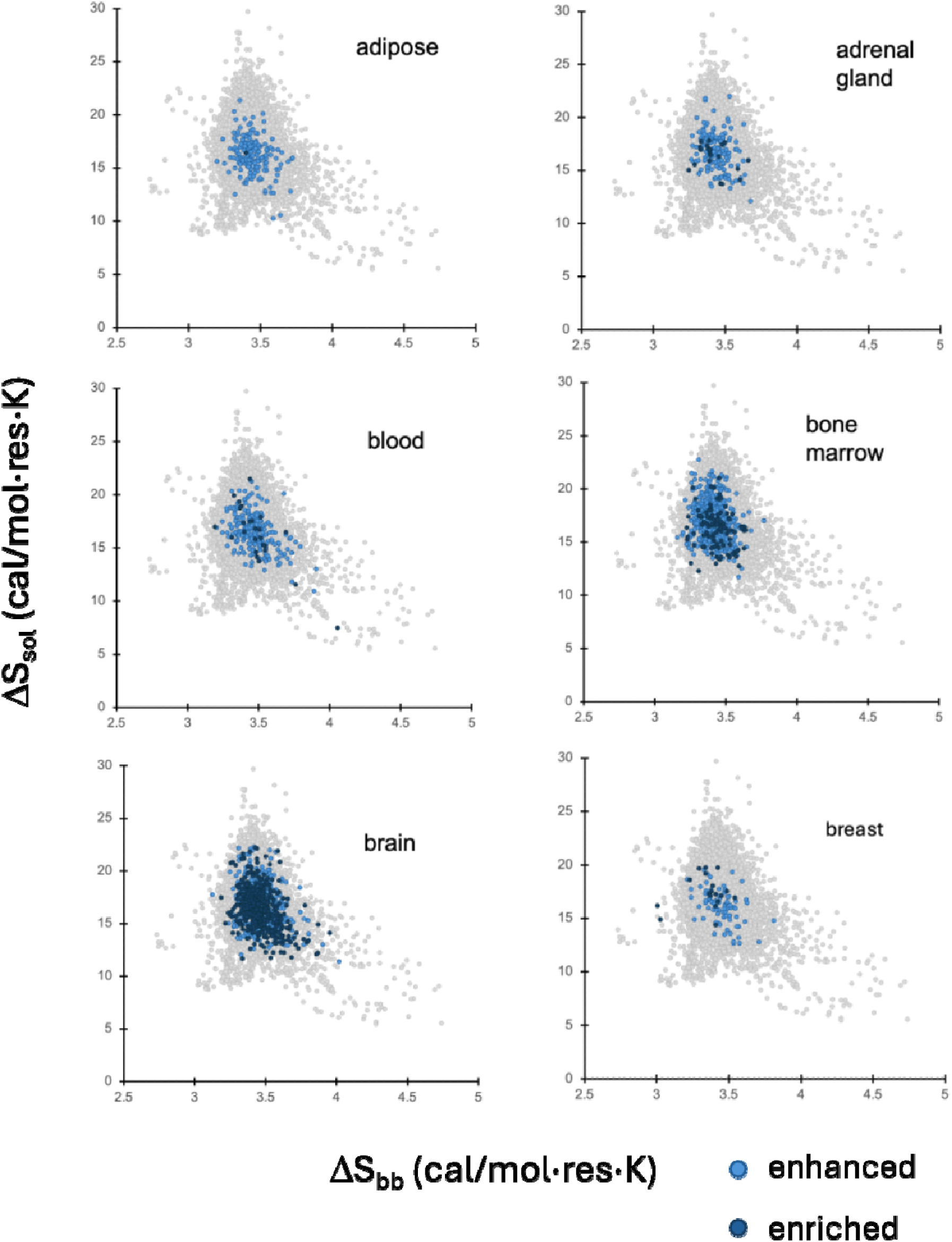
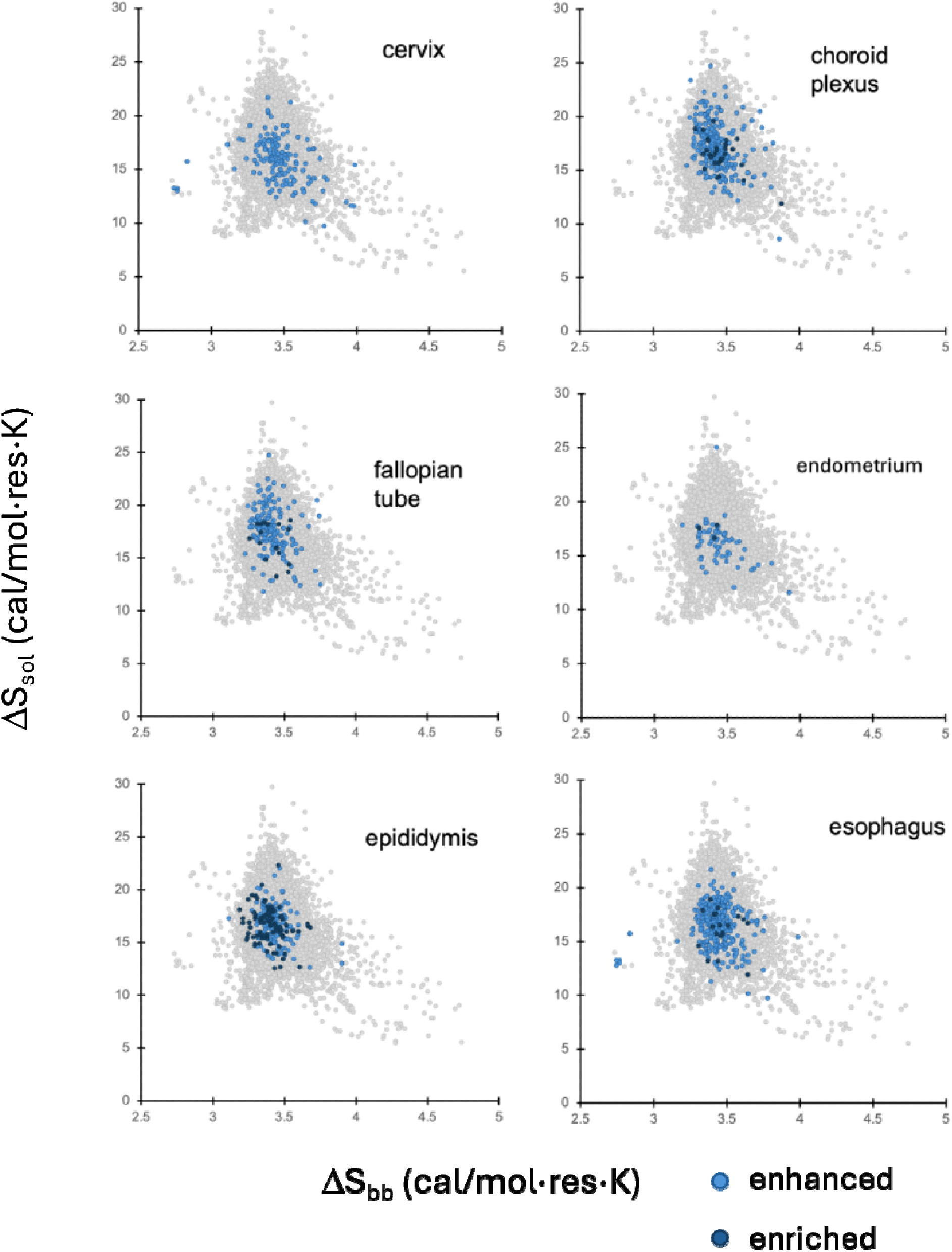
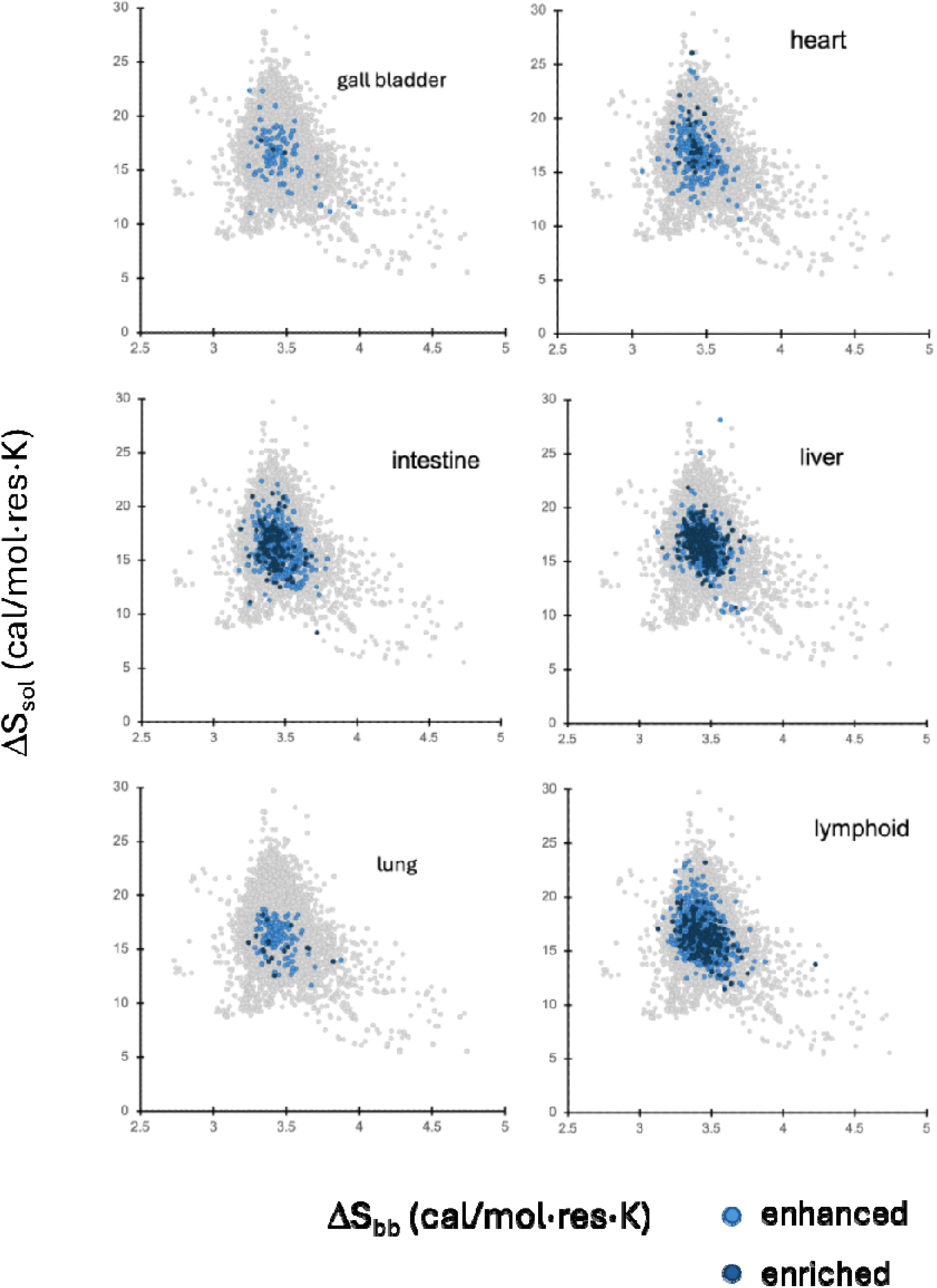
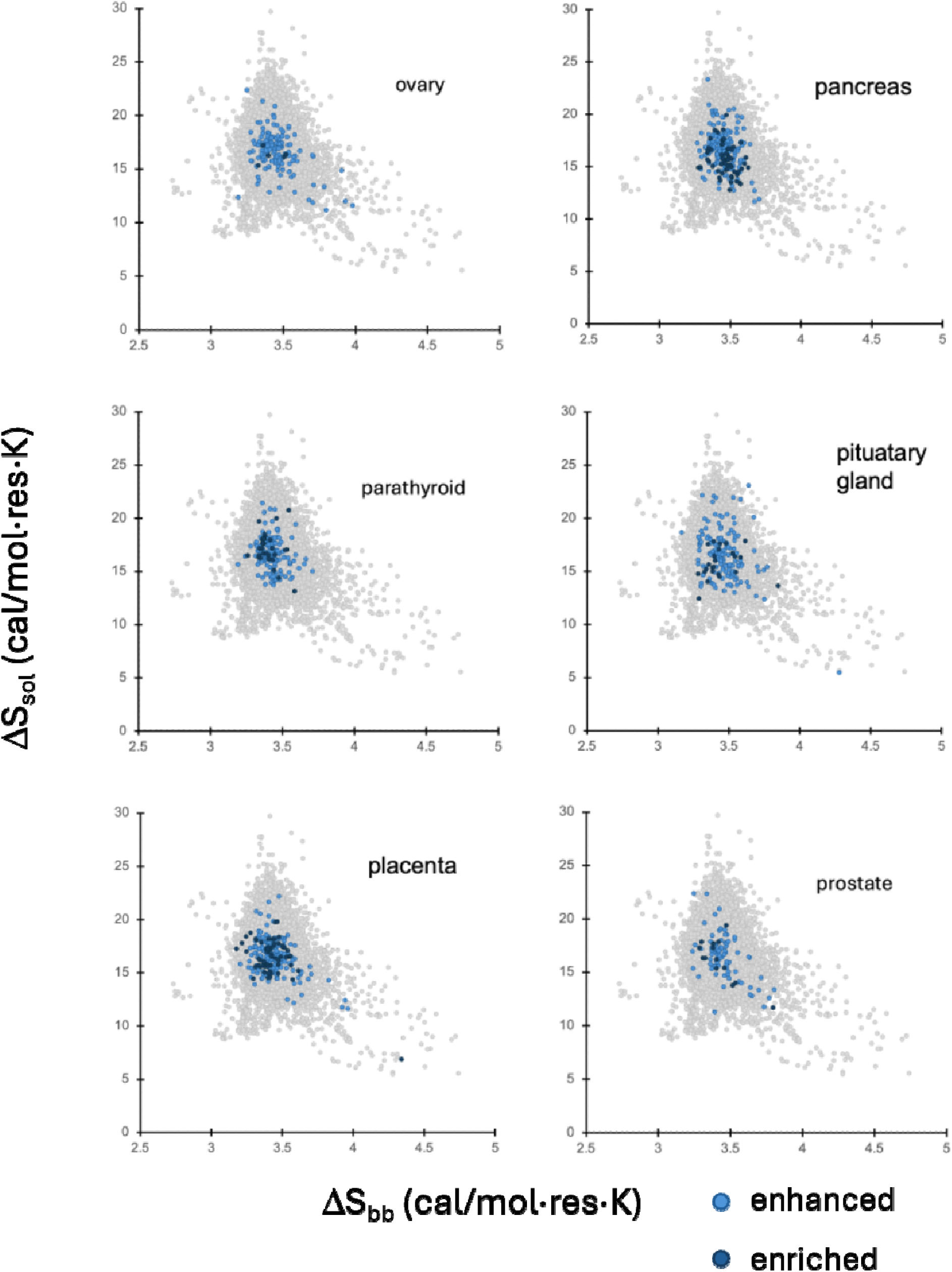
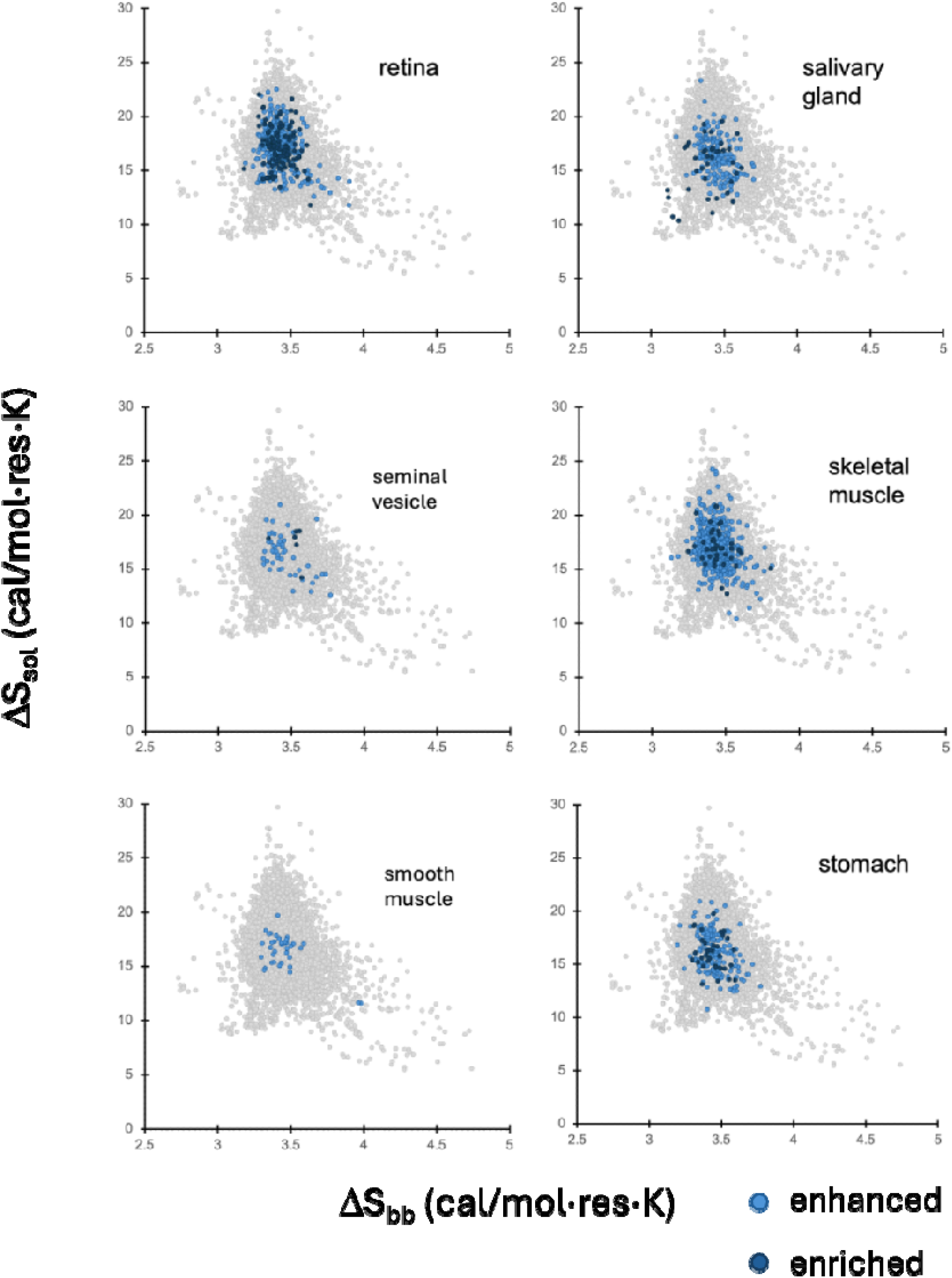
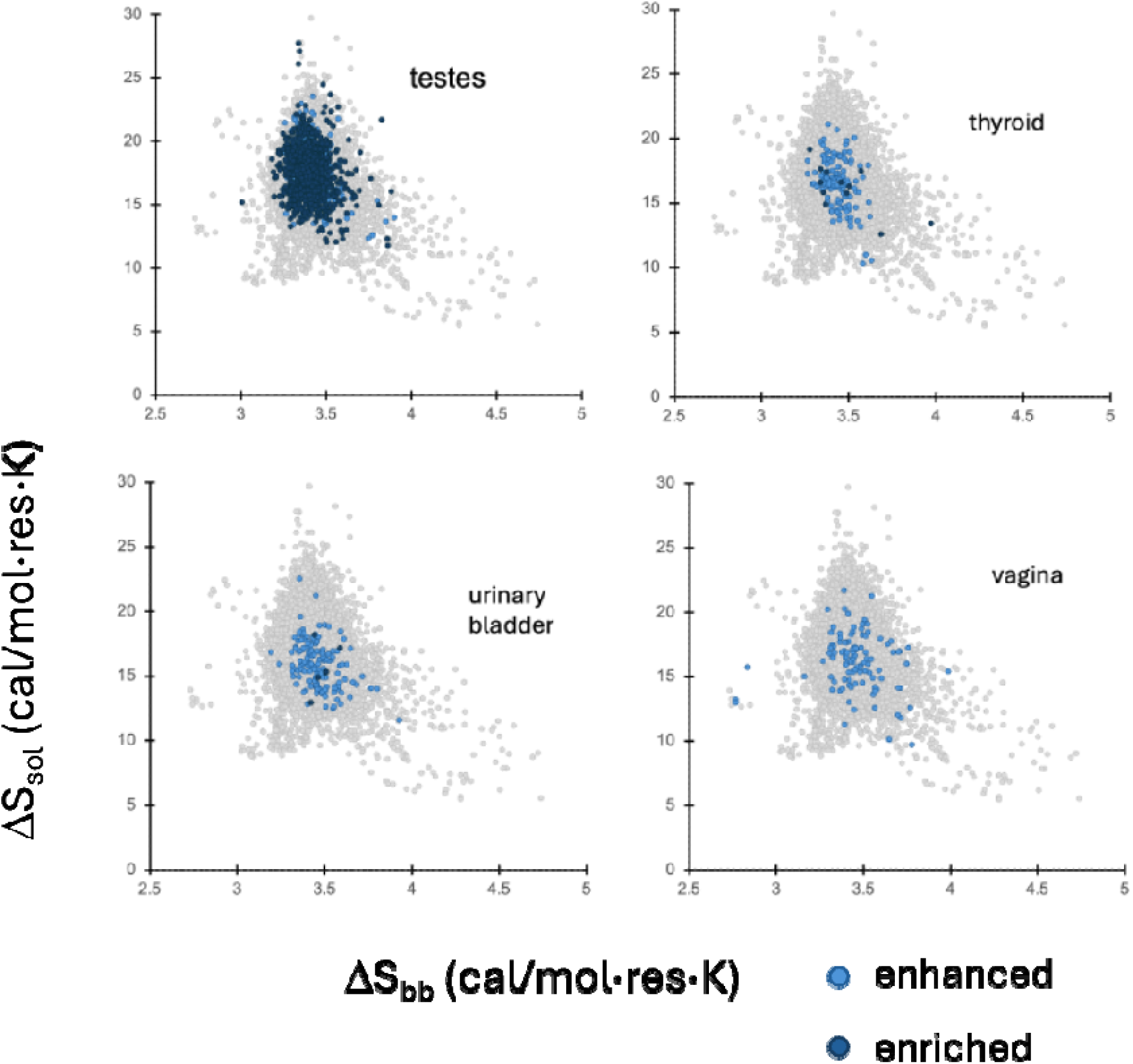
Tissue-level transcriptome entropy maps based on the Human Protein Atlas. [41]. Skin and kidney maps are in Fig. 4.

**Figure S3.**
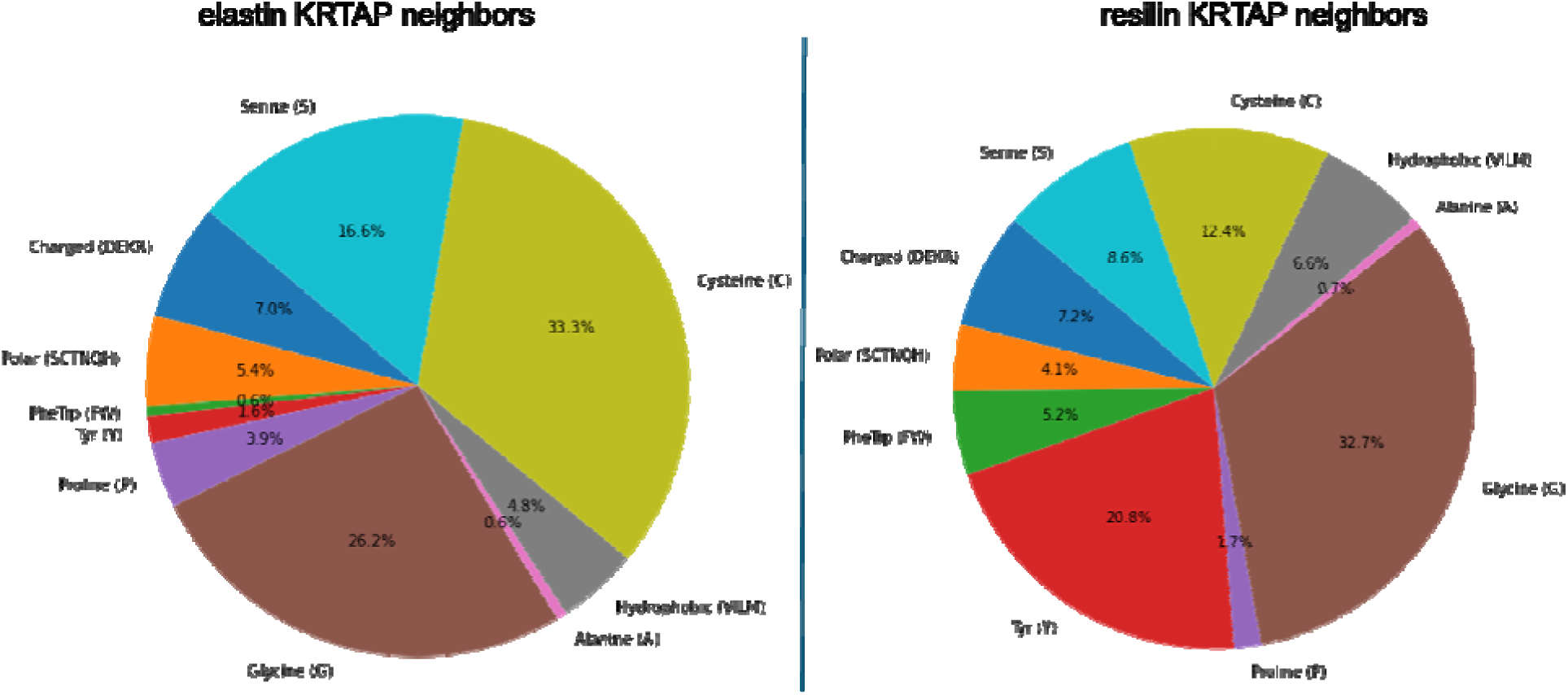
Amino acid compositions of the keratin-associated proteins. (KRTAPs) in the elastin or resilin whole protein neighborhood.

**Figure S4.**
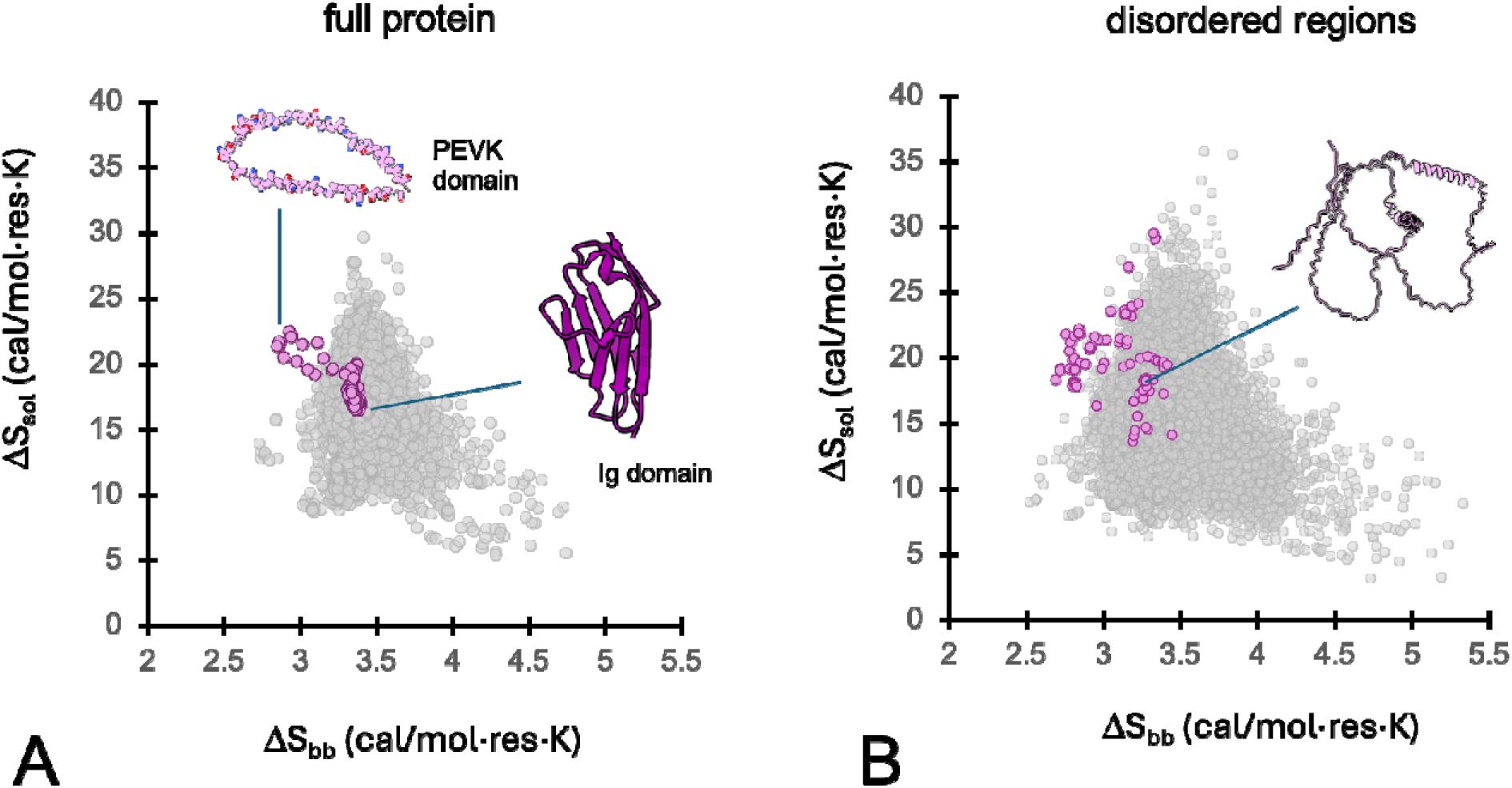
PEVK domain of titin in the whole protein and disordered domain entropy maps.

**Figure S5.**
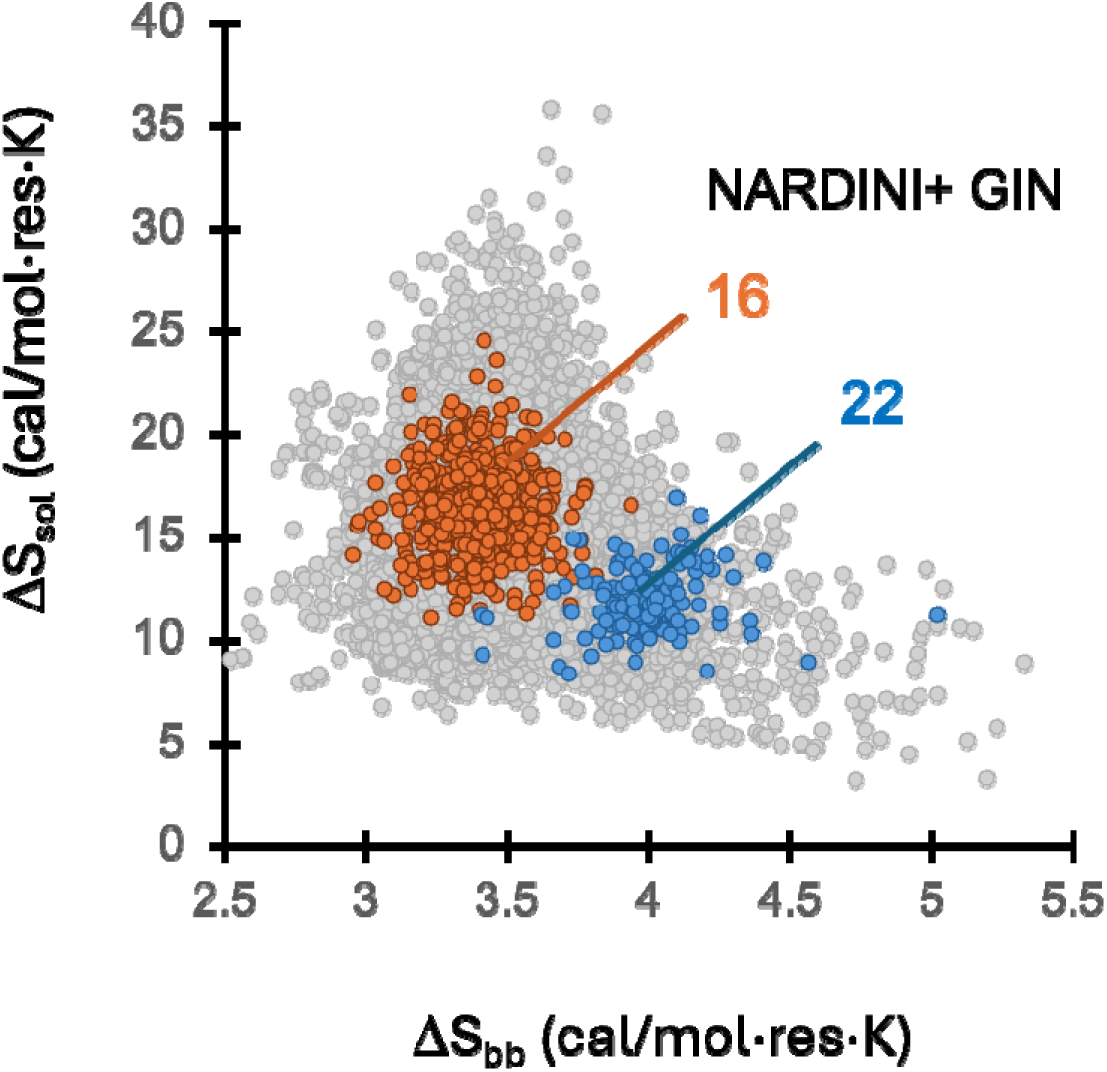
Sequence grammar classes of elastic proteins. Disordered protein sequences were projected onto the grammar-based NARDINI+ GIN classification framework of Ruff, Pappu and colleagues [4], which groups intrinsically disordered regions according to residue composition and sequence patterning. Cluster 22, identified as an elastomer-enriched class, is characterized by well-mixed Gly-and Pro-rich sequences and maps predominantly to the high ΔS_bb_, low ΔS_sol_ region of the entropy landscape occupied by resilin and many candidate disorder-based elastic proteins. A fragment of elastin exon 26 maps to Cluster 16, a grammar class enriched in blocky distributions of polar residues that occupies a more central position in entropy space.

**Figure S6.**
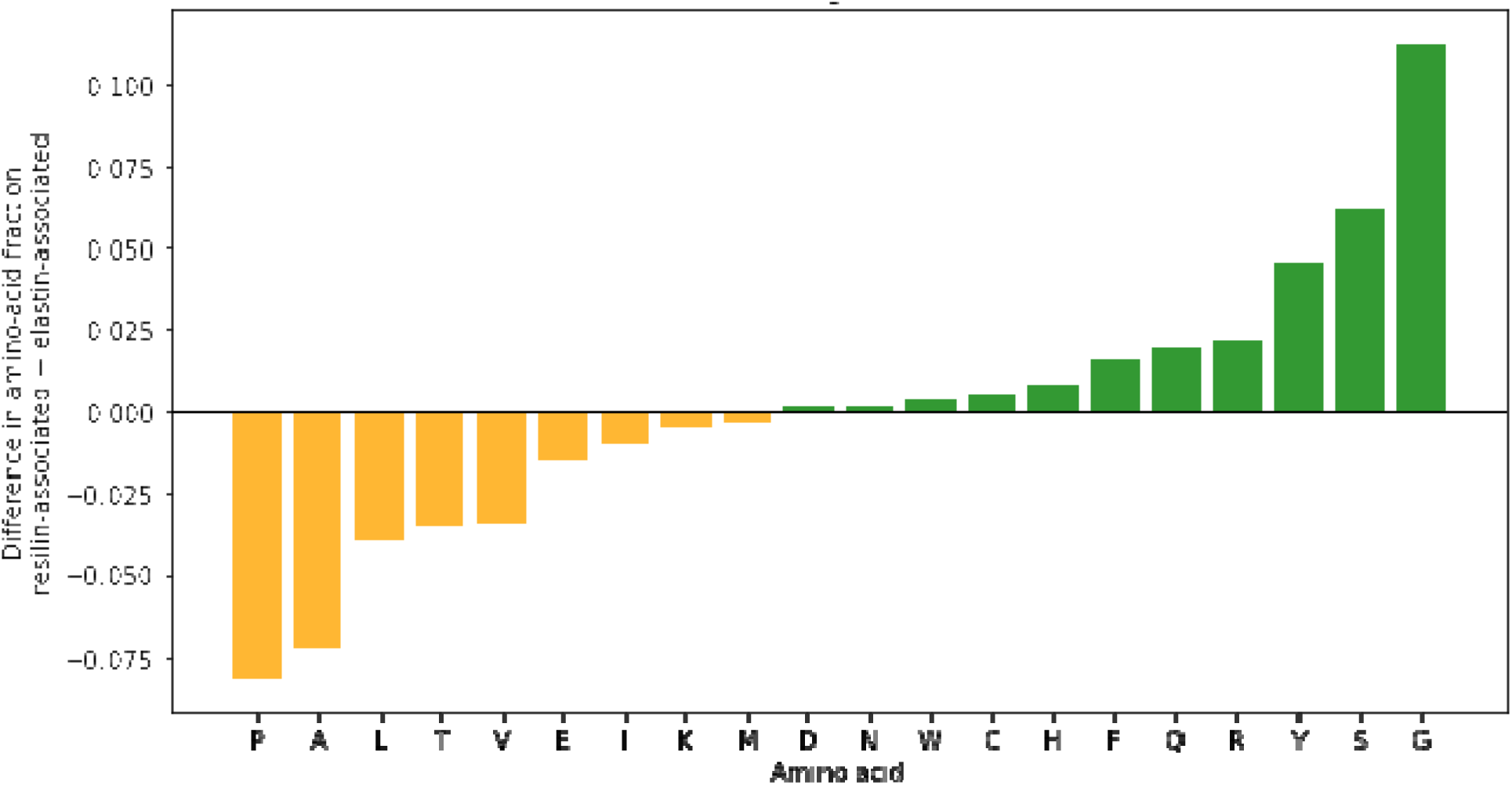
Amino acid compositional biases distinguishing elastin-and resilin-associated protein language model neighborhoods. Relative amino acid enrichment was calculated by comparing the aggregate amino acid frequencies of proteins in the resilin PLM kNN neighborhood with those in the elastin PLM kNN neighborhood. Positive values (green) indicate residues enriched in the resilin-associated network, negative values (orange) indicate residues enriched in the elastin-associated network.

## SUPPLEMENTARY TABLES

**Table S1.** Backbone and solvation entropy scales.

| | $\Delta S_{bb}^a$<br>(cal/mol·K) | $-T\Delta S^b$<br>(kcal/mol) | $\Delta S_{sol}^c$<br>(cal/mol·K) |
| --- | --- | --- | --- |
| **P** | 1.8<sup>d</sup> | 16 | 6.7 |
| **V** | 2.18 | 17.2 | 10.7 |
| **I** | 2.18 | 18.1 | 13.7 |
| **C** | 3.4 | 15.3 | 4.3 |
| **S** | 3.4 | 15.6 | 5.3 |
| **T** | 3.4 | 16.7 | 9.0 |
| **L** | 3.4 | 17.8 | 12.7 |
| **M** | 3.4 | 18.1 | 13.7 |
| **N** | 3.4 | 19.1 | 17.0 |
| **F** | 3.4 | 19.9 | 19.7 |
| **Y** | 3.4 | 20.2 | 20.7 |
| **Q** | 3.4 | 20.6 | 22.0 |
| **H** | 3.4 | 21.1 | 23.7 |
| **W** | 3.4 | 22.3 | 27.7 |
| **K** | 3.4 | 22.8 | 29.3 |
| **R** | 3.4 | 24.1 | 33.7 |
| **D** | 3.4 | 26.4 | 41.3 |
| **E** | 3.4 | 27.7 | 45.7 |
| **A** | 4.1 | 14.8 | 2.7 |
| **G** | 6.5 | 14 | 0.0 |

<sup>a</sup> values from D'Aquino et. al. Table II [16], <sup>b</sup> values from Schauperl et. al. Table I for TIP3P water [23], <sup>c</sup>calculated assuming $T = 300K$ and baseline normalized for glycine, <sup>d</sup>estimated assuming fraction of Ramachandran space occupied for proline and glycine $F_{Pro}/F_{Gly} = 0.1$ .

**Table S2.**
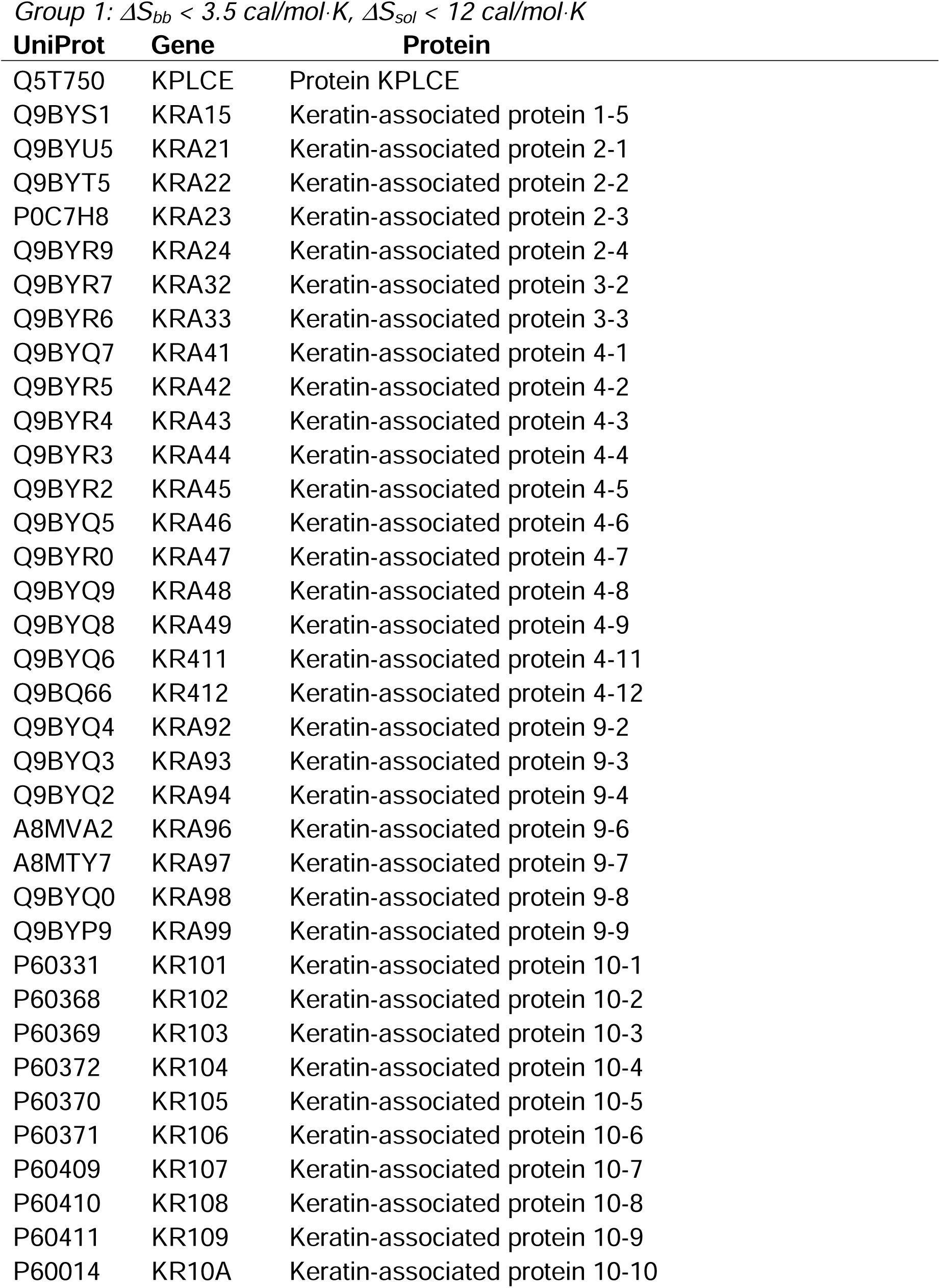
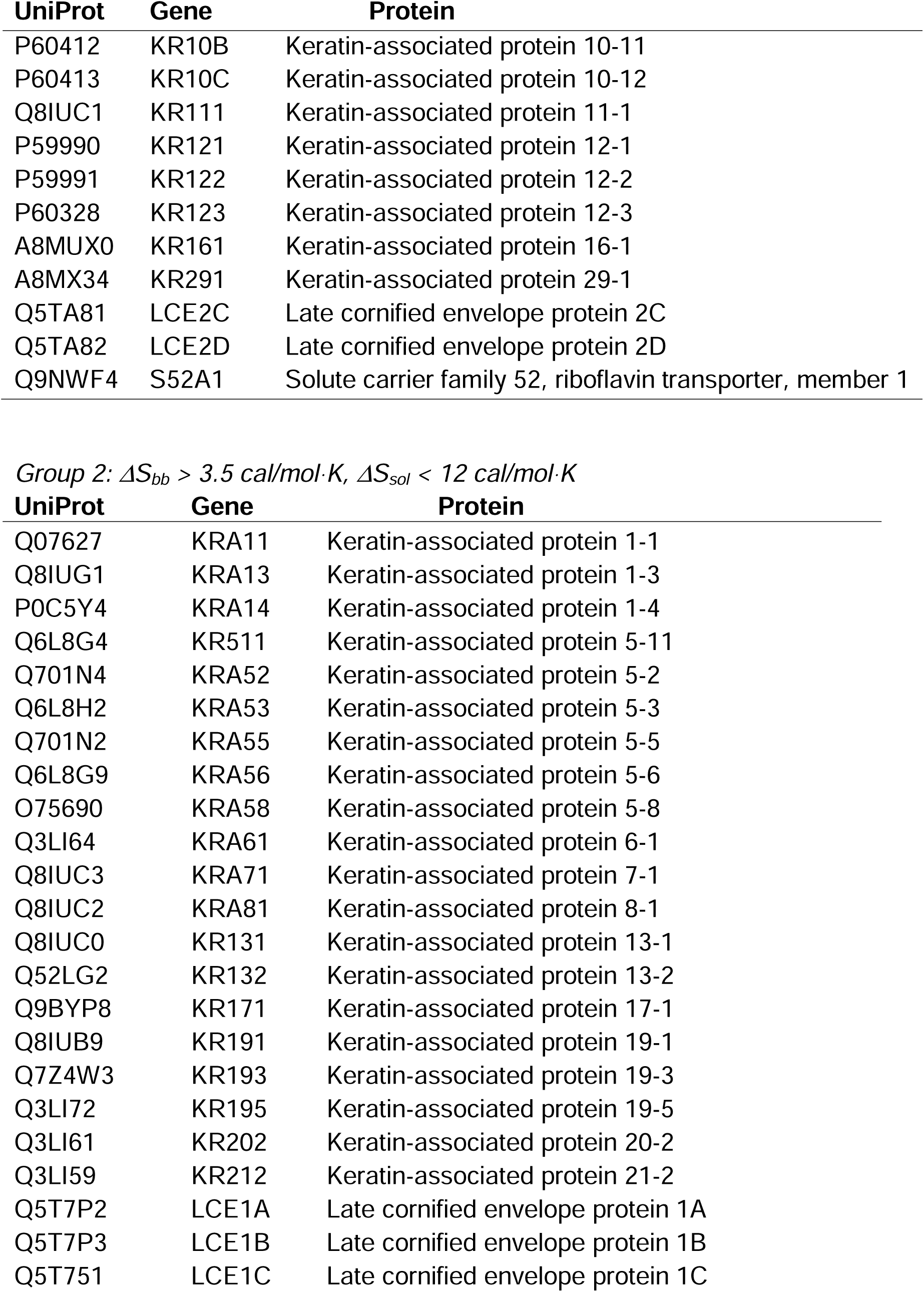
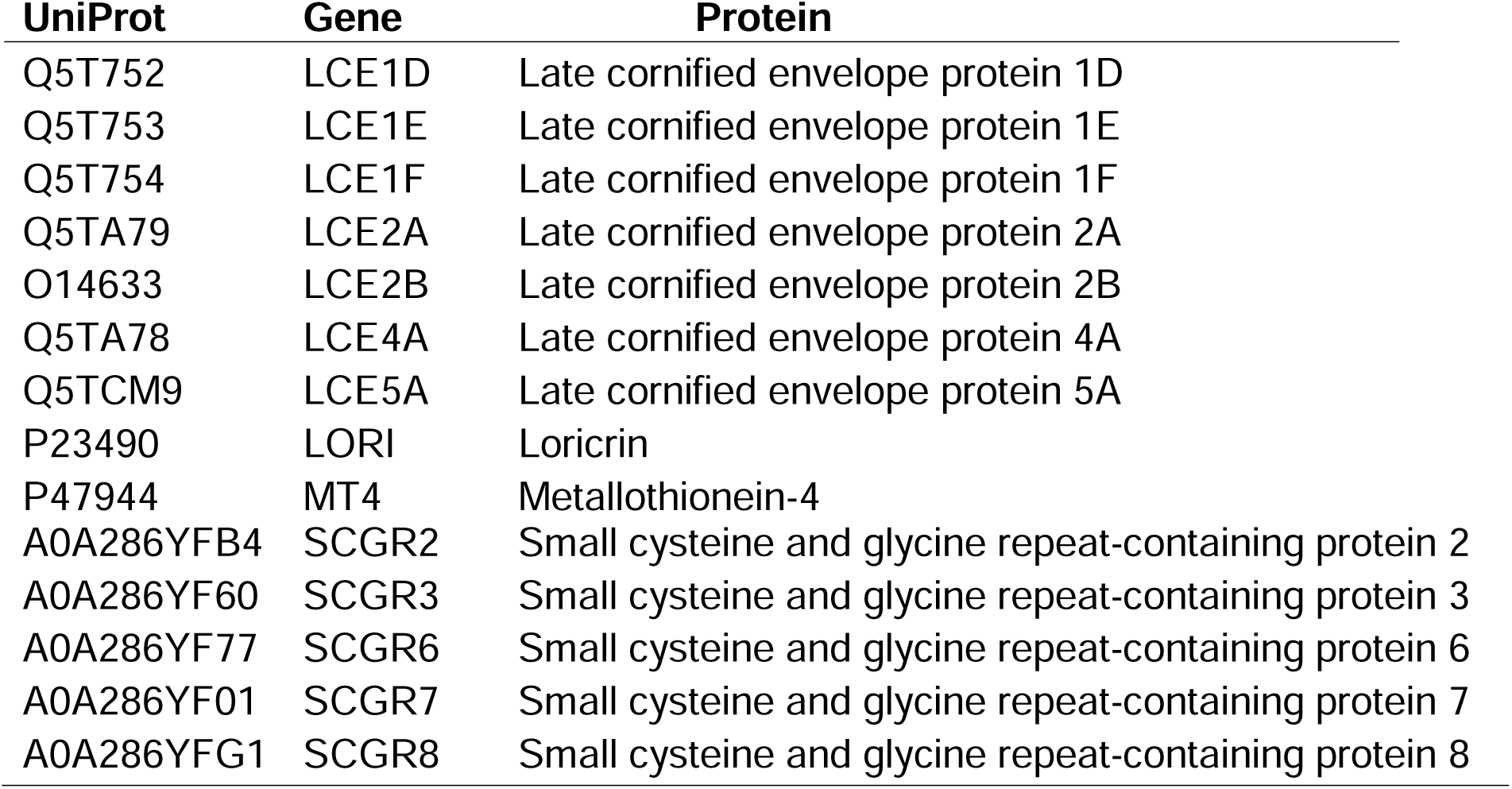
Skin enriched proteins – whole protein sequence (see Fig. 3A)

**Table S3.**
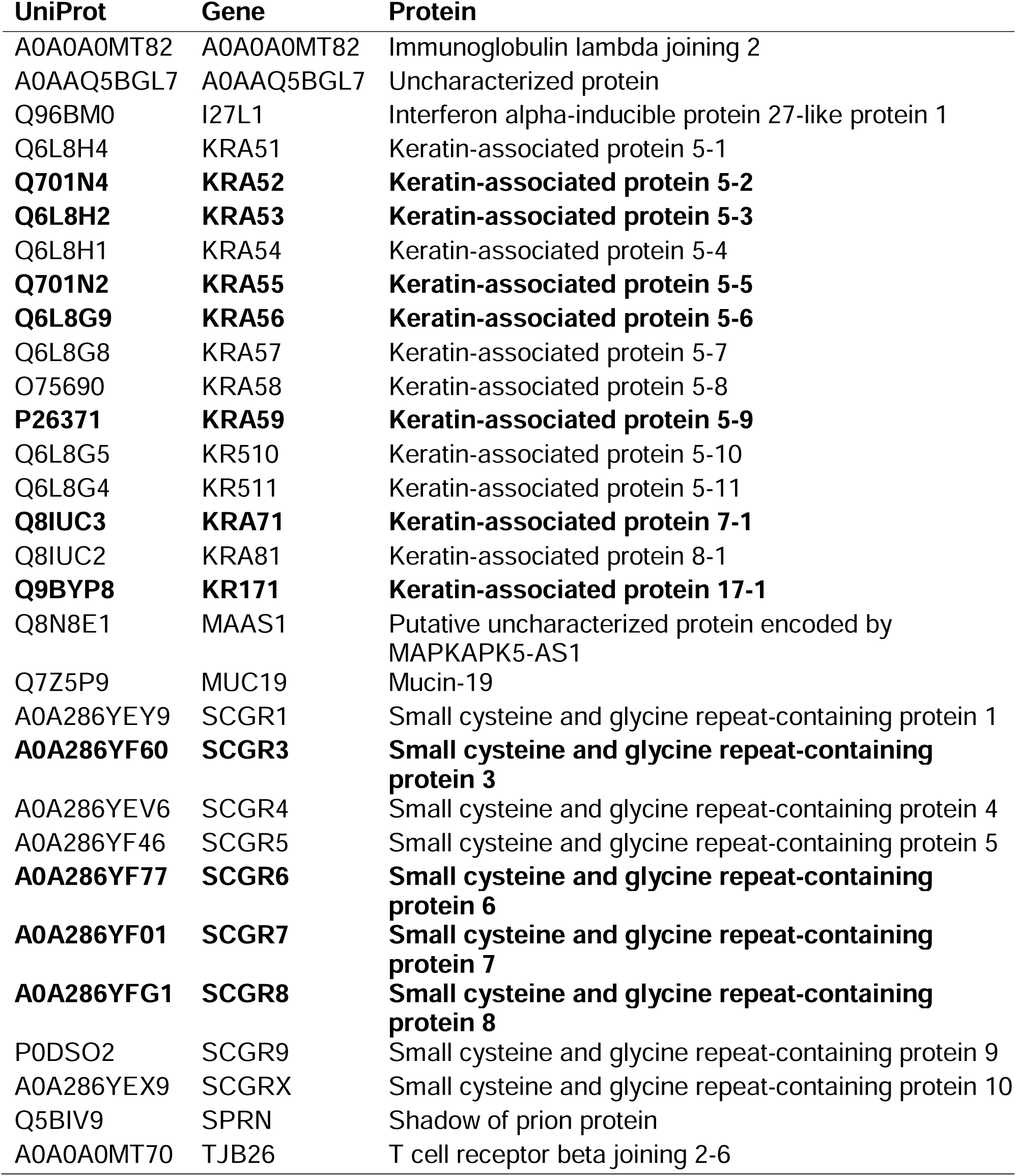
Elastin neighbors – whole protein sequences (from Fig. 2A). Skin enriched proteins in bold.

**Table S4.**
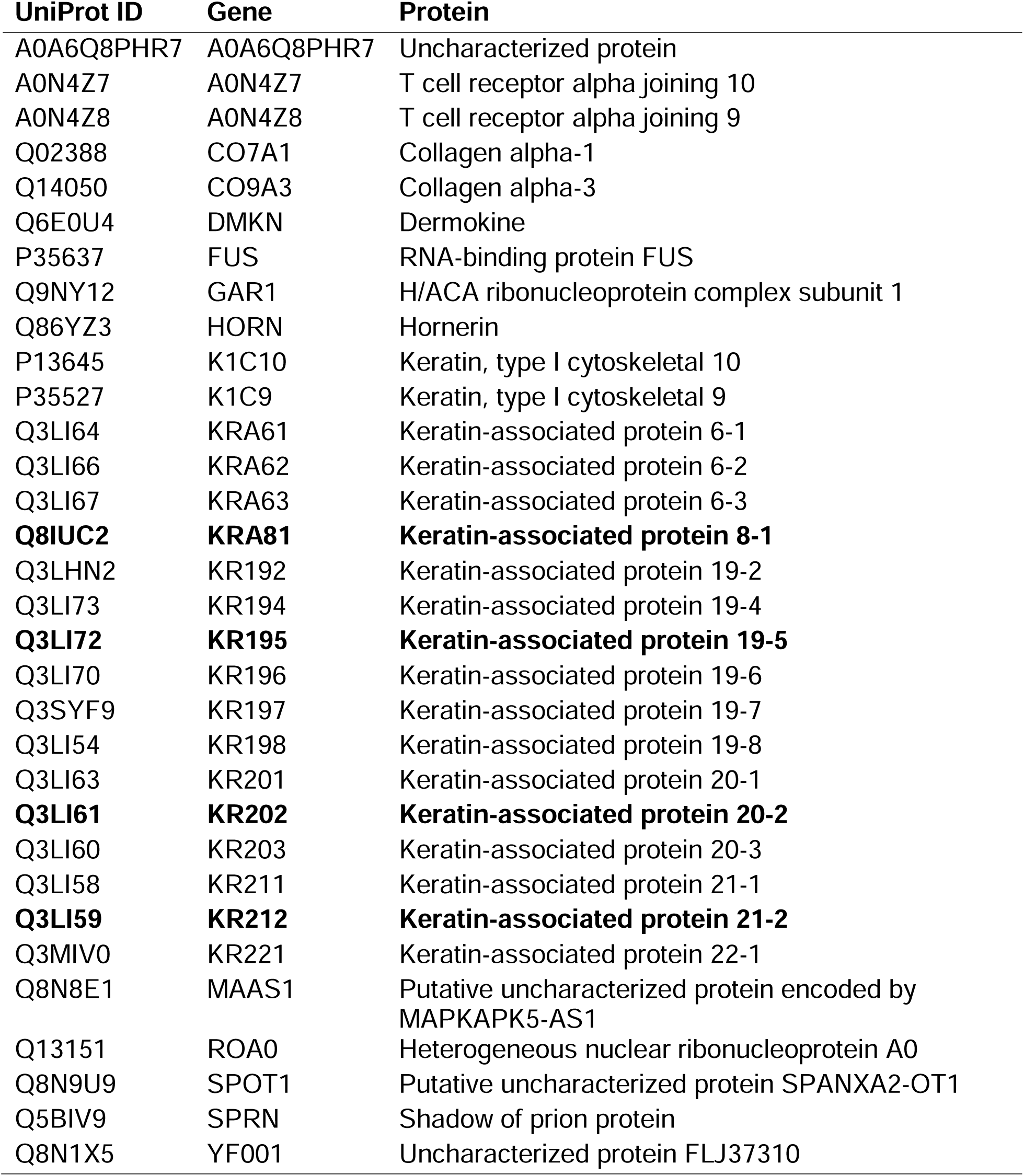
Resilin neighbors – whole protein sequences (from Fig. 2A). Skin enriched proteins in bold.

| UniProt ID | Gene | Protein |
| --- | --- | --- |
| A0A6Q8PHR7 | A0A6Q8PHR7 | Uncharacterized protein |
| A0N4Z7 | A0N4Z7 | T cell receptor alpha joining 10 |
| A0N4Z8 | A0N4Z8 | T cell receptor alpha joining 9 |
| Q02388 | CO7A1 | Collagen alpha-1 |
| Q14050 | CO9A3 | Collagen alpha-3 |
| Q6E0U4 | DMKN | Dermokine |
| P35637 | FUS | RNA-binding protein FUS |
| Q9NY12 | GAR1 | H/ACA ribonucleoprotein complex subunit 1 |
| Q86YZ3 | HORN | Hornerin |
| P13645 | K1C10 | Keratin, type I cytoskeletal 10 |
| P35527 | K1C9 | Keratin, type I cytoskeletal 9 |
| Q3LI64 | KRA61 | Keratin-associated protein 6-1 |
| Q3LI66 | KRA62 | Keratin-associated protein 6-2 |
| Q3LI67 | KRA63 | Keratin-associated protein 6-3 |
| **Q8IUC2** | **KRA81** | **Keratin-associated protein 8-1** |
| Q3LHN2 | KR192 | Keratin-associated protein 19-2 |
| Q3LI73 | KR194 | Keratin-associated protein 19-4 |
| **Q3LI72** | **KR195** | **Keratin-associated protein 19-5** |
| Q3LI70 | KR196 | Keratin-associated protein 19-6 |
| Q3SYF9 | KR197 | Keratin-associated protein 19-7 |
| Q3LI54 | KR198 | Keratin-associated protein 19-8 |
| Q3LI63 | KR201 | Keratin-associated protein 20-1 |
| **Q3LI61** | **KR202** | **Keratin-associated protein 20-2** |
| Q3LI60 | KR203 | Keratin-associated protein 20-3 |
| Q3LI58 | KR211 | Keratin-associated protein 21-1 |
| **Q3LI59** | **KR212** | **Keratin-associated protein 21-2** |
| Q3MIV0 | KR221 | Keratin-associated protein 22-1 |
| Q8N8E1 | MAAS1 | Putative uncharacterized protein encoded by MAPKAPK5-AS1 |
| Q13151 | ROA0 | Heterogeneous nuclear ribonucleoprotein A0 |
| Q8N9U9 | SPOT1 | Putative uncharacterized protein SPANXA2-OT1 |
| Q5BIV9 | SPRN | Shadow of prion protein |
| Q8N1X5 | YF001 | Uncharacterized protein FLJ37310 |

**Table S5.** Elastin neighbors – disordered domains. (from Fig. 5B)

| UniProt | Start | Stop | Length | Gene | Protein |
| --- | --- | --- | --- | --- | --- |
| O75912 | 1 | 107 | 1057 | DGKI | Diacylglycerol kinase iota |
| O95678 | 463 | 551 | 551 | K2C75 | Keratin, type II cytoskeletal 75 |
| P02538 | 1 | 153 | 564 | K2C6A | Keratin, type II cytoskeletal 6A |
| P02538 | 480 | 564 | 564 | K2C6A | Keratin, type II cytoskeletal 6A |
| P04259 | 1 | 153 | 564 | K2C6B | Keratin, type II cytoskeletal 6B |
| P04259 | 480 | 564 | 564 | K2C6B | Keratin, type II cytoskeletal 6B |
| P08727 | 1 | 75 | 400 | K1C19 | Keratin, type I cytoskeletal 19 |
| P0DTL6 | 1 | 76 | 404 | ZTRF1 | Zinc finger TRAF-type-containing protein 1 |
| P19013 | 452 | 520 | 520 | K2C4 | Keratin, type II cytoskeletal 4 |
| P20264 | 1 | 102 | 500 | PO3F3 | POU domain, class 3, transcription factor 3 |
| P31314 | 1 | 168 | 330 | TLX1 | T-cell leukemia homeobox protein 1 |
| P35556 | 18 | 101 | 2912 | FBN2 | Fibrillin-2 [Cleaved into: Placensin] |
| P35556 | 220 | 301 | 2912 | FBN2 | Fibrillin-2 [Cleaved into: Placensin] |
| P35556 | 419 | 501 | 2912 | FBN2 | Fibrillin-2 [Cleaved into: Placensin] |
| P41225 | 64 | 131 | 446 | SOX3 | Transcription factor SOX-3 |
| P48668 | 1 | 153 | 564 | K2C6C | Keratin, type II cytoskeletal 6C |
| P48668 | 480 | 564 | 564 | K2C6C | Keratin, type II cytoskeletal 6C |
| P51513 | 371 | 421 | 507 | NOVA1 | RNA-binding protein Nova-1 |
| P52948 | 1 | 157 | 1817 | NUP98 | Nuclear pore complex protein Nup98-Nup96 |
| P56270 | 214 | 277 | 477 | MAZ | Myc-associated zinc finger protein |
| P84550 | 272 | 470 | 965 | SKOR1 | SKI family transcriptional corepressor 1 |
| Q00653 | 341 | 435 | 900 | NFKB2 | Nuclear factor NF-kappa-B p100 subunit |
| Q01546 | 498 | 638 | 638 | K22O | Keratin, type II cytoskeletal 2 oral |
| Q01851 | 72 | 225 | 419 | PO4F1 | POU domain, class 4, transcription factor 1 |
| Q12816 | 796 | 1431 | 1431 | TROP | Trophinin |
| Q12948 | 318 | 486 | 553 | FOXC1 | Forkhead box protein C1 |
| Q14050 | 68 | 149 | 684 | CO9A3 | Collagen alpha-3 |
| Q6P1L5 | 41 | 139 | 589 | F117B | Protein FAM117B |
| Q7Z3B4 | 1 | 111 | 507 | NUP54 | Nucleoporin p54 |
| Q7Z3Y9 | 1 | 80 | 468 | K1C26 | Keratin, type I cytoskeletal 26 |
| Q7Z3Z0 | 1 | 76 | 450 | K1C25 | Keratin, type I cytoskeletal 25 |
| Q7Z4P5 | 19 | 82 | 450 | GDF7 | Growth/differentiation factor 7 |
| Q7Z5P9 | 108 | 357 | 8384 | MUC19 | Mucin-19 |
| Q7Z5P9 | 150 | 650 | 8384 | MUC19 | Mucin-19 |
| Q7Z5P9 | 581 | 849 | 8384 | MUC19 | Mucin-19 |
| Q7Z5P9 | 1204 | 1275 | 8384 | MUC19 | Mucin-19 |
| Q86XN8 | 1 | 51 | 651 | MEX3D | RNA-binding protein MEX3D |
| Q8IVW8 | 34 | 93 | 549 | SPNS2 | Sphingosine-1-phosphate transporter<br>SPNS2 |
| Q8NFJ8 | 38 | 227 | 381 | BHE22 | Class E basic helix-loop-helix protein 22 |
| Q8NFP9 | 1 | 68 | 2946 | NBEA | Neurobeachin |
| Q96F45 | 1 | 61 | 646 | ZN503 | Zinc finger protein 503 |
| Q9BXB5 | 1 | 76 | 764 | OSB10 | Oxysterol-binding protein-related protein 10 |
| Q9H0L4 | 489 | 570 | 616 | CSTFT | Cleavage stimulation factor subunit 2 tau<br>variant |
| Q9H7D7 | 1 | 118 | 661 | WDR26 | WD repeat-containing protein 26 |
| Q9HCR9 | 46 | 117 | 933 | PDE11 | Dual 3',5'-cyclic-AMP and -GMP<br>phosphodiesterase 11A |
| Q9UBV8 | 19 | 109 | 284 | PEF1 | Peflin |
| Q9Y2X9 | 1 | 90 | 895 | ZN281 | Zinc finger protein 281 |
| Q9Y4X0 | 1 | 122 | 333 | AMMR1 | Nuclear protein AMMECR1 |

**Table S6.**
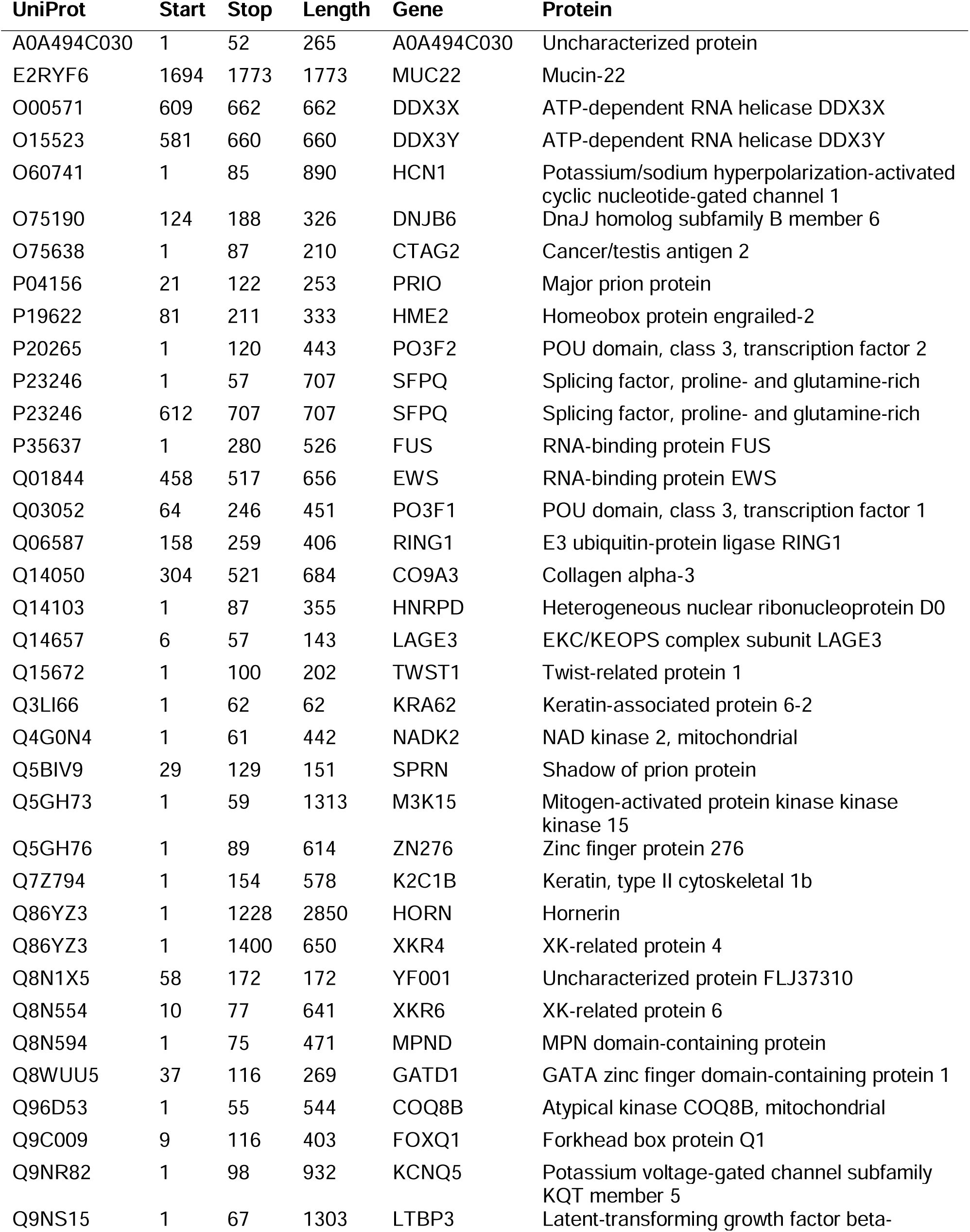
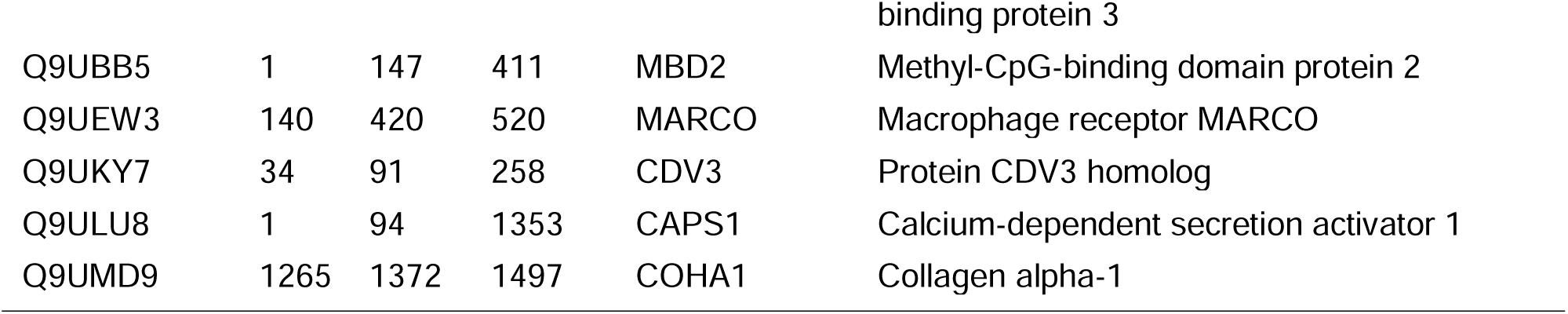
Resilin neighbors – disordered domains. (from Fig. 5B)

| UniProt | Start | Stop | Length | Gene | Protein |
| --- | --- | --- | --- | --- | --- |
| A0A494C030 | 1 | 52 | 265 | A0A494C030 | Uncharacterized protein |
| E2RYF6 | 1694 | 1773 | 1773 | MUC22 | Mucin-22 |
| O00571 | 609 | 662 | 662 | DDX3X | ATP-dependent RNA helicase DDX3X |
| O15523 | 581 | 660 | 660 | DDX3Y | ATP-dependent RNA helicase DDX3Y |
| O60741 | 1 | 85 | 890 | HCN1 | Potassium/sodium hyperpolarization-activated cyclic nucleotide-gated channel 1 |
| O75190 | 124 | 188 | 326 | DNJB6 | DnaJ homolog subfamily B member 6 |
| O75638 | 1 | 87 | 210 | CTAG2 | Cancer/testis antigen 2 |
| P04156 | 21 | 122 | 253 | PRIO | Major prion protein |
| P19622 | 81 | 211 | 333 | HME2 | Homeobox protein engrailed-2 |
| P20265 | 1 | 120 | 443 | PO3F2 | POU domain, class 3, transcription factor 2 |
| P23246 | 1 | 57 | 707 | SFPQ | Splicing factor, proline- and glutamine-rich |
| P23246 | 612 | 707 | 707 | SFPQ | Splicing factor, proline- and glutamine-rich |
| P35637 | 1 | 280 | 526 | FUS | RNA-binding protein FUS |
| Q01844 | 458 | 517 | 656 | EWS | RNA-binding protein EWS |
| Q03052 | 64 | 246 | 451 | PO3F1 | POU domain, class 3, transcription factor 1 |
| Q06587 | 158 | 259 | 406 | RING1 | E3 ubiquitin-protein ligase RING1 |
| Q14050 | 304 | 521 | 684 | CO9A3 | Collagen alpha-3 |
| Q14103 | 1 | 87 | 355 | HNRPD | Heterogeneous nuclear ribonucleoprotein D0 |
| Q14657 | 6 | 57 | 143 | LAGE3 | EKC/KEOPS complex subunit LAGE3 |
| Q15672 | 1 | 100 | 202 | TWST1 | Twist-related protein 1 |
| Q3LI66 | 1 | 62 | 62 | KRA62 | Keratin-associated protein 6-2 |
| Q4G0N4 | 1 | 61 | 442 | NADK2 | NAD kinase 2, mitochondrial |
| Q5BIV9 | 29 | 129 | 151 | SPRN | Shadow of prion protein |
| Q5GH73 | 1 | 59 | 1313 | M3K15 | Mitogen-activated protein kinase kinase kinase 15 |
| Q5GH76 | 1 | 89 | 614 | ZN276 | Zinc finger protein 276 |
| Q7Z794 | 1 | 154 | 578 | K2C1B | Keratin, type II cytoskeletal 1b |
| Q86YZ3 | 1 | 1228 | 2850 | HORN | Hornerin |
| Q86YZ3 | 1 | 1400 | 650 | XKR4 | XK-related protein 4 |
| Q8N1X5 | 58 | 172 | 172 | YF001 | Uncharacterized protein FLJ37310 |
| Q8N554 | 10 | 77 | 641 | XKR6 | XK-related protein 6 |
| Q8N594 | 1 | 75 | 471 | MPND | MPN domain-containing protein |
| Q8WUU5 | 37 | 116 | 269 | GATD1 | GATA zinc finger domain-containing protein 1 |
| Q96D53 | 1 | 55 | 544 | COQ8B | Atypical kinase COQ8B, mitochondrial |
| Q9C009 | 9 | 116 | 403 | FOXQ1 | Forkhead box protein Q1 |
| Q9NR82 | 1 | 98 | 932 | KCNQ5 | Potassium voltage-gated channel subfamily KQT member 5 |
| Q9NS15 | 1 | 67 | 1303 | LTBP3 | Latent-transforming growth factor beta- binding protein 3 |
| Q9UBB5 | 1 | 147 | 411 | MBD2 | Methyl-CpG-binding domain protein 2 |
| Q9UEW3 | 140 | 420 | 520 | MARCO | Macrophage receptor MARCO |
| Q9UKY7 | 34 | 91 | 258 | CDV3 | Protein CDV3 homolog |
| Q9ULU8 | 1 | 94 | 1353 | CAPS1 | Calcium-dependent secretion activator 1 |
| Q9UMD9 | 1265 | 1372 | 1497 | COHA1 | Collagen alpha-1 |

**Table S7.**
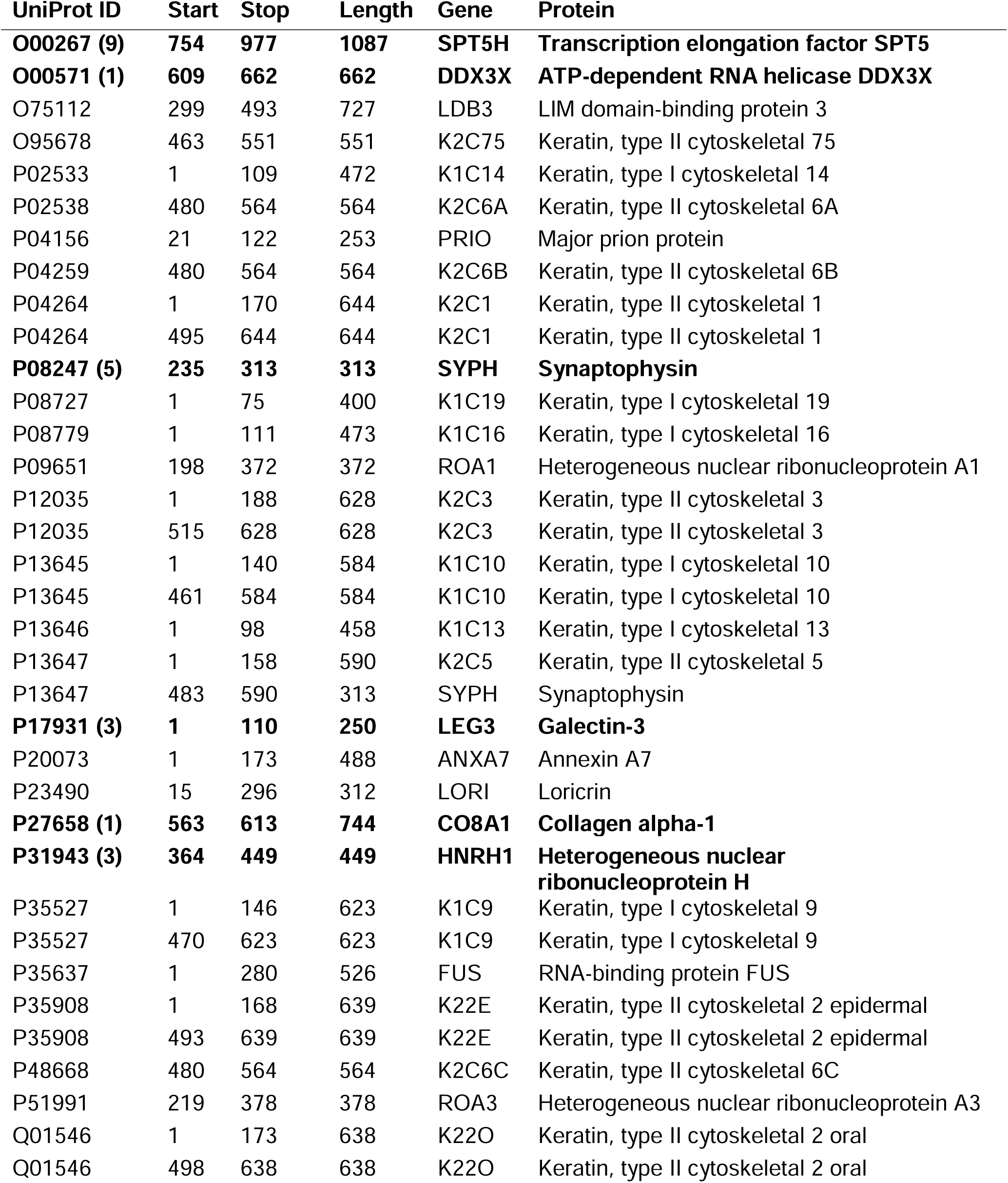
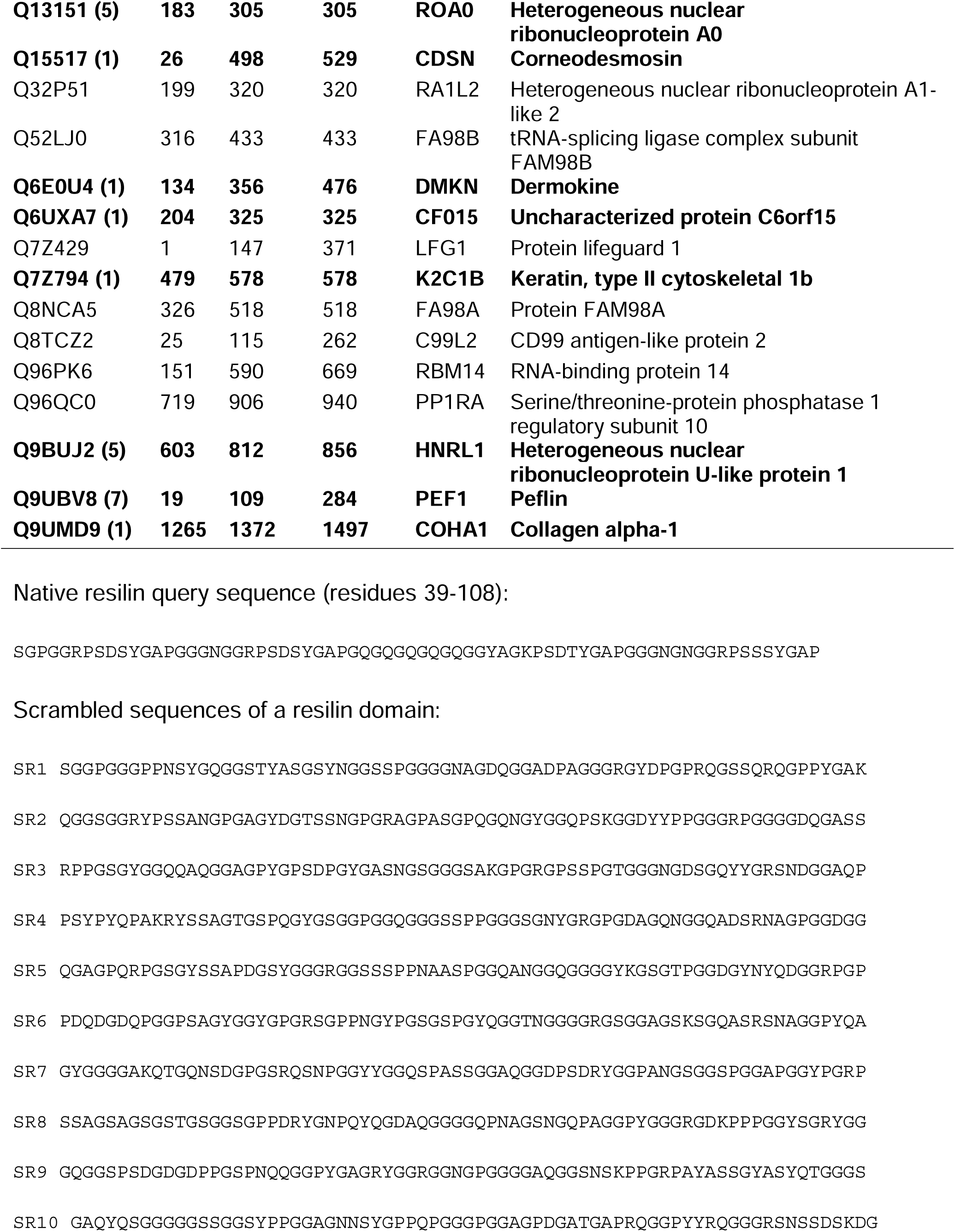
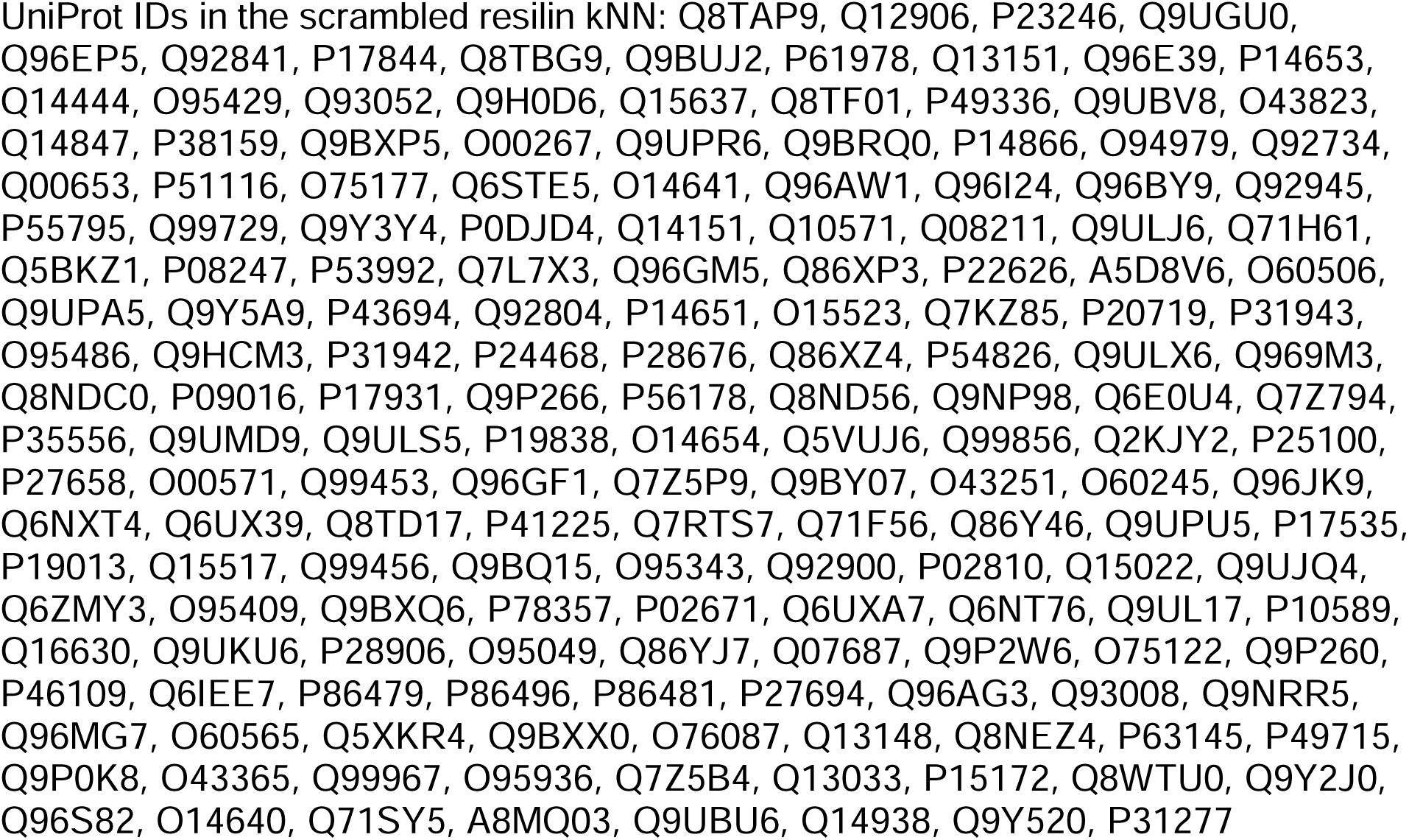
Resilin kNN in the PLM – disordered domains. Bolded entries also appears in the scrambled sequence search with number of matches in parentheses.

**Table S8.**
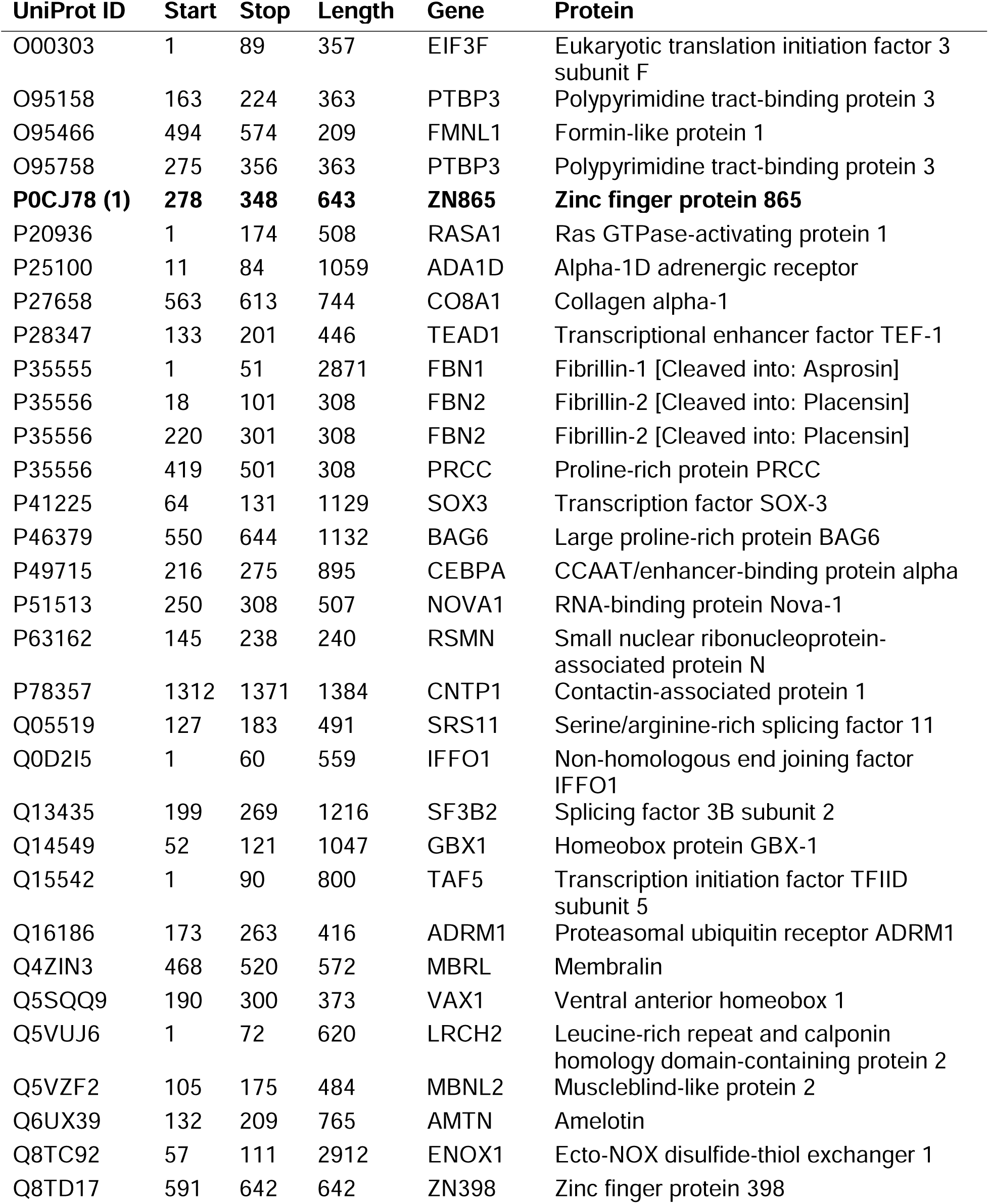
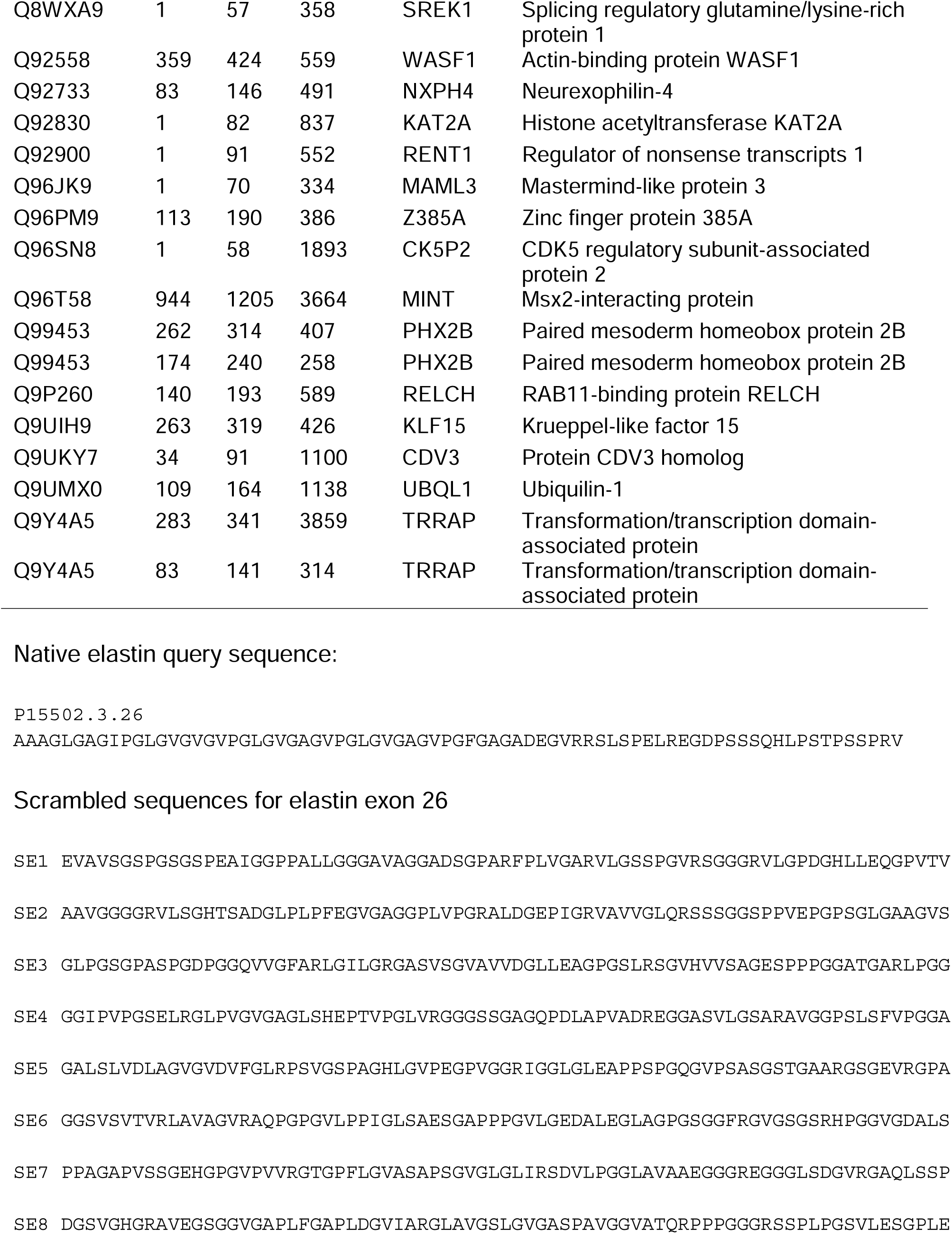
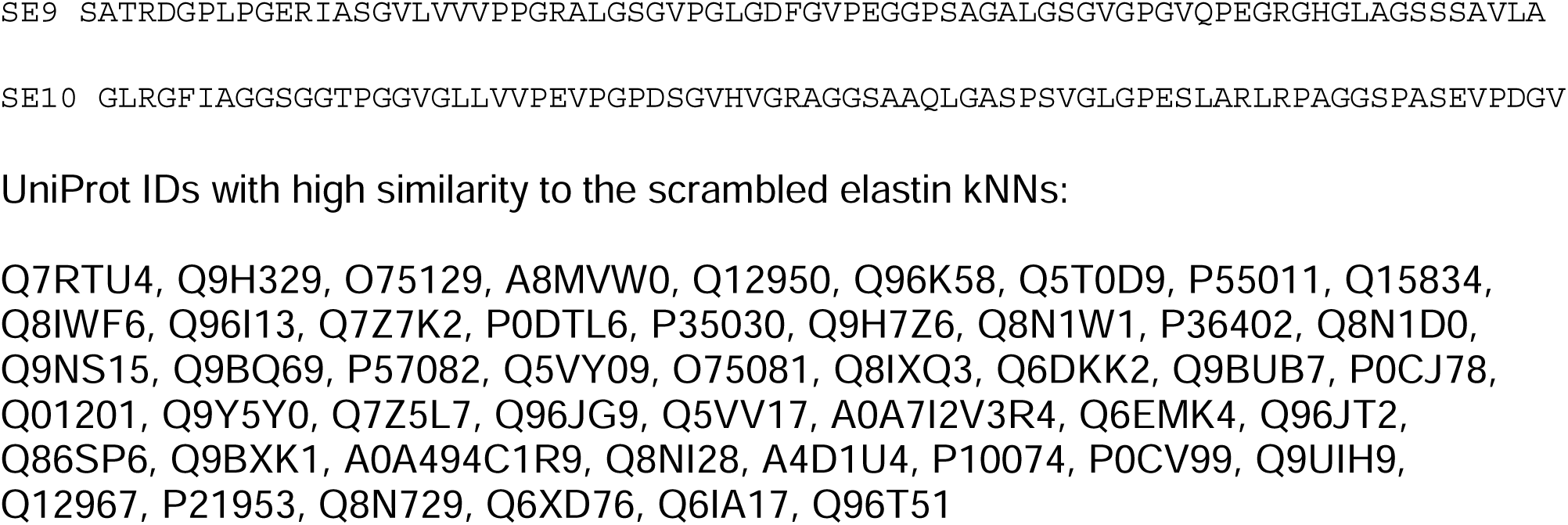
Elastin kNN in the PLM – disordered domains. Bolded entry also appears in scrambled sequence search with number of matches in parentheses.

